# Bioinformatic Analysis of Structure and Function of LIM Domains of Human Zyxin Family Proteins

**DOI:** 10.1101/2020.07.14.202572

**Authors:** Mohammad Quadir Siddiqui, Maulik D. Badmalia, Trushar R. Patel

## Abstract

Members of the human Zyxin family are LIM domain-containing proteins that perform critical cellular functions and are indispensable for cellular integrity. Despite their importance, not much is known about their structure, functions, interactions and dynamics. To provide insights into these, we used a set of in-silico tools and databases and analyzed their amino acid sequence, phylogeny, post-translational modifications, structure-dynamics, molecular interactions, and functions. Our analysis revealed that zyxin members are ohnologs. Presence of a conserved nuclear export signal composed of LxxLxL/LxxxLxL consensus sequence, as well as a possible nuclear localization signal, suggesting that Zyxin family members may have nuclear and cytoplasmic roles. The molecular modeling and structural analysis indicated that Zyxin family LIM domains share similarities with transcriptional regulators and have positively charged electrostatic patches, which may indicate that they have previously unanticipated nucleic acid binding properties. Intrinsic dynamics analysis of Lim domains suggest that only Lim1 has similar internal dynamics properties, unlike Lim2/3. Furthermore, we analyzed protein expression and mutational frequency in various malignancies, as well as mapped protein-protein interaction networks they are involved in. Overall, our comprehensive bioinformatic analysis suggests that these proteins may play important roles in mediating protein-protein and protein-nucleic acid interactions.

## 1. Introduction

Zyxin family of proteins is a family of LIM-domain containing proteins that play crucial roles in mediating cellular signalling, tumorigenesis and developmental pathways, and have, therefore, been implicated in human disease [1–5]. For example, Zyxin family proteins have been reported to be involved in various cancers including prostate, colorectal, lung, liver and melanoma [6–9]. Moreover, the founding member of this family, zyxin has recently been identified as a potential biomarker for the aggressive phenotypes of human brain cancer (glioblastoma multiforme), exhibits differential expression levels in melanocytes/melanoma cells and altered phosphorylation in colon cancer, and is found to be associated with poor prognosis in glioma patients [10].

In addition to zyxin, other family members have also been linked to the cancer predisposition (see **Table 1** for a full list of Zyxin family members). For example, WTIP has a crucial role in cell proliferation and downregulation of WTIP is associated with poor prognosis and survival of non-small-cell lung cancer patients [11]. Interestingly, LIMD1, Ajuba, and WTIP were discovered as novel mammalian processing body (P-body) components and implicated in novel mechanisms of miRNA-mediated gene silencing [12]. FBLIM1 promotes cell migration and invasion in human glioma by altering the PLC-γ/STAT3 signaling pathway and could be used as a molecular marker for early diagnosis in glioma patients [13]. LPP was required for TGF-β induced cell migration/adhesion dynamics and regulates the invadopodia formation with SHCA adapter protein cooperation [14], and loss of LPP/Etv5/MMP-15 may be implicated in the prognostic marker of lung adenocarcinoma [15].

**Table 1:** List of Zyxin family proteins containing Lim domain, functions and biochemical properties.

| Protein<br>(UNIPROT<br>ID) | Cellular role * | Cellular<br>localization | Isoelectric point<sup>1</sup> |  | Amino acids /<br>Molecular weight (kDa)<sup>1</sup> |  |
| --- | --- | --- | --- | --- | --- | --- |
|  |  |  | Full<br>protein | LIM<br>domain | Full pro-<br>tein | LIM domain |
| Zyxin<br>(Q15942) | Adhesion plague<br>protein, signal<br>transducer | Cytoskeleton,<br>cytosol and nu-<br>cleus | 6.22 | 7.16 | 572/<br>61.27 | 189/21.10 |
| LimD1<br>(Q9UGP4) | Cell fate determi-<br>nation and a tu-<br>mour suppressor | Predominantly<br>in nucleus, P-body | 6.20 | 5.90 | 676/<br>72.19 | 195/21.87 |

|  |  |  |  |  |  |  |
| --- | --- | --- | --- | --- | --- | --- |
| Ajuba<br>(Q96IF1) | Cell fate determination and repression of gene transcription | Cell membrane, cytoskeleton, centrosome, nucleus, P- body | 6.86 | 5.81 | 538/<br>56.93 | 195/21.87 |
| LPP<br>(Q93052) | Maintains cell shape, motility, and activation of gene transcription | Cell membrane and nucleus | 7.18 | 6.78 | 612/<br>65.74 | 190/21.15 |
| WTIP<br>(A6NIX2) | Cell fate determination, repression of gene transcription, and miRNA-mediated gene silencing | Nucleus, P- body | 8.53 | 7.50 | 430/<br>45.12 | 195/21.80 |
| FBLIM1<br>(Q8WUP2) | Cell-ECM adhesion proteins and filamin-containing actin filaments | Focal adhesion sites | 5.71 | 7.79 | 373/<br>40.66 | 190/21.70 |
| TRIP6<br>(Q15654) | Relays signals from the cell surface to the nucleus | Focal adhesion sites, cytoplasm, nucleus | 7.19 | 6.06 | 476/<br>50.28 | 188/20.49 |

\*= Information retrieved from UniprotKB
<sup>1</sup>= Calculated using ProtParam

Although members of the Zyxin family proteins are functionally diverse, playing important roles from cytoskeleton to transcriptional machinery, their domain organization is relatively conserved. They are characterized by the presence of the N-terminal proline-rich region (PRR) and three LIM (*Lin-11, Isl-1, and Mec-3*) domains, LIM1-3 at their C-terminal region **(Figure 1**). In general, it is believed that Zyxin family of proteins execute their diverse functions by interacting with cellular proteins via their PRR and Lim domains that serve as the docking sites for binding [1, 16–18]. However, despite their functional relevance and importance for human disease physiology, many details about the structure, functions and dynamics of Zyxin family members are still missing, thus significantly limiting their utility as key targets. In addition, not much is known about how relatively similar architecture translates into such a wide range of functions.

**Figure 1:**
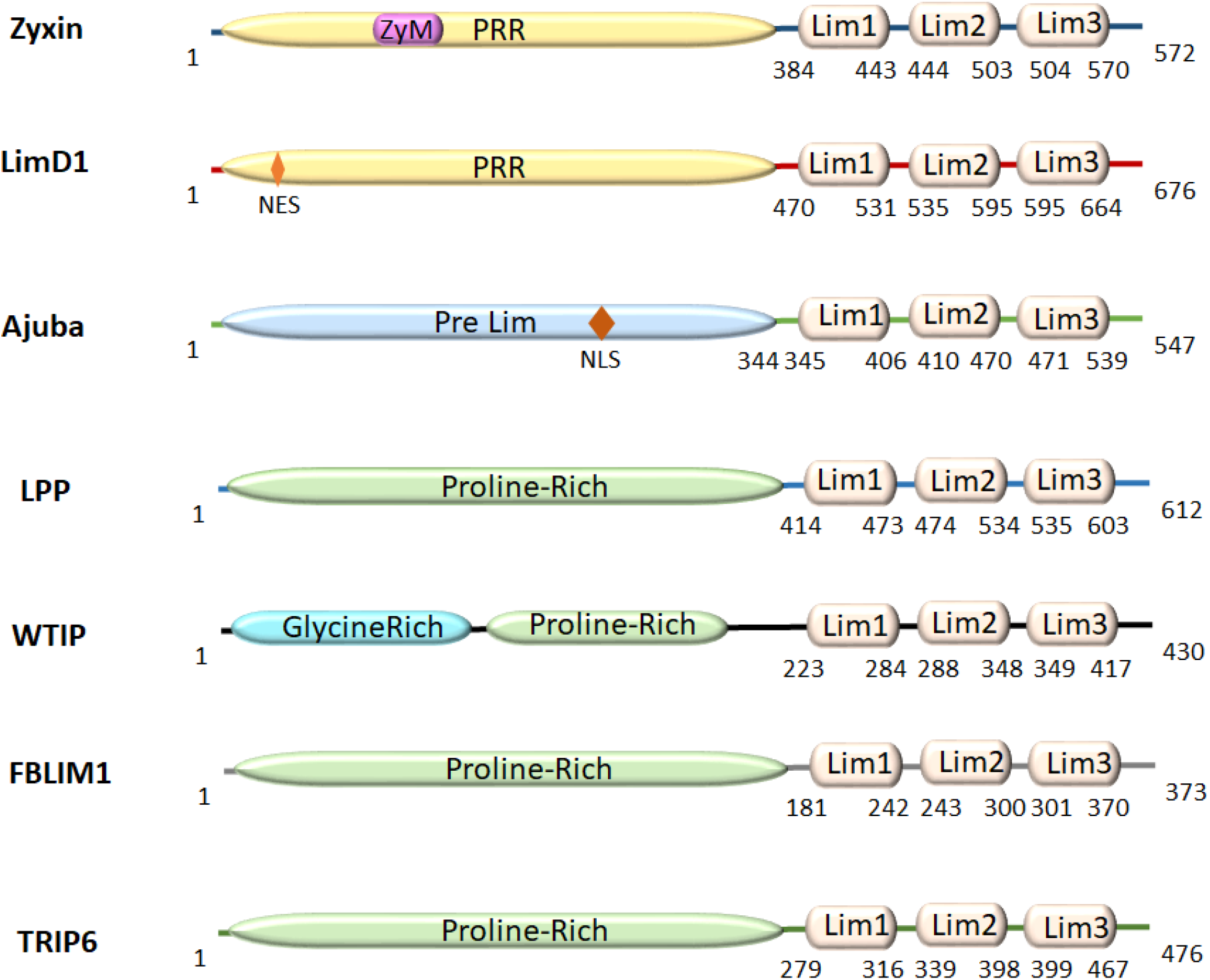
(A) Organization of domains and motifs of Zyxin family proteins: the proline-rich region (PRR) and Lim1/2/3 domains.

Here, we address this knowledge gap and provide insights into fundamental questions related to the structure and function of human Zyxin family members. We used various bioinformatics tools to dissect sequence features, phylogeny, post-translational modifications, structure-dynamics, molecular interactions, and functions of these proteins. Our analysis reveals that these proteins are ohnologs and have a conserved nuclear export sequence (NES). Two members also suggests the presence of a potential nuclear localization sequence (NLS), therefore supporting the notion that these proteins shuttle in the cell nucleus. Furthermore, structural homologous models of LIM1-3 regions suggest a high similarity with transcription regulators. Analysis of the electrostatic surface potential and DNAbinder further support that these LIM domains may engage in nucleic acid-binding. Based on our investigation of structure-dynamic features we propose that differences in protein structure/dynamics, may serve as the key determinants of functional variability in this family. Additionally, analyses of protein expression and mutational frequency in various malignancies, as well as the Zyxin family of protein interactomes, indicate that these proteins may contribute to cancer by regulating metabolic rewiring and oncogene addiction. Collectively, our bioinformatic analysis has important implications for follow-up functional and structural investigations of Zyxin family proteins.

## 2. Results

### 2.1. Human Zyxin family proteins exhibit variations in the proline-rich region and Lim edges

The amino acid sequence composition of Zyxin family proteins varies considerably. Interestingly, our sequence alignment showed that within Lim domains, the presence of cysteine/histidine/aspartate residues and their spacing is quite uniform **(Figures 1 & Supplementary Figure 1A),** and follows the LIM domain consensus sequence (Kadrmas and Beckerle 2004). We observed that, as expected, each LIM domain contains a set of four cysteine/histidine residues that in other LIM domains are known to coordinate Zn^2+^ ions leading to the formation of Zinc finger topology arranged in tandem [18, 19]. At the sequence level, we observed the presence of three distinct LIM domains (LIM1-3). Each LIM domain exhibited a high level of conservation among proposed Zn^2+^ion coordinating residues, as well as the spacing between these, suggesting a high degree of sequence conservation among Lim domains in members of the Zyxin family (Zyxin, LimD1, Ajuba, LPP, WTIP, FBLIM1, TRIP6) **(Supplementary Figure 1A).** Moreover, each LIM domain was found to be ~55-65 amino acids long with each finger consisting of ~30 amino acids. Additionally, the sequence analysis revealed that the N-terminal PRR, which is devoid of cysteine/histidine residues, contains repeated unit clusters of 2 to 6 amino acids not seen in the C-terminal LIM domains. The overall sequence length of these Zyxin family members varies substantially from 373 to 676 amino acids, with PRR constituting ~48-69%, and LIM domains occupying ~30-51% of the polypeptide chains **(Table 2).**

**Table 2:** Percentage composition of Proline Rich Region (PRR) and Lim domain in the full-length proteins

| Protein<br>(Uniprot ID) | Percentage of PRR vs.<br>Full length* | Percentage of Lim domain<br>vs. Full length* |
| --- | --- | --- |
| Zyxin (Q15942) | 66.9% | 33.1% |
| LimD1 (Q9UGP4) | 69.3% | 30.7% |
| Ajuba (Q96IF1) | 62.8% | 37.2% |
| LPP (Q93052) | 67.4% | 32.6% |
| WTIP (A6NIX2) | 51.6% | 48.4% |
| FBLIM1<br>(Q8WUP2) | 48.2% | 51.8% |
| TRIP6 (Q15654) | 58.4% | 41.6% |

\*= Calculated by considering full length (all amino acids) as a 100% of the respective protein PRR= Proline Rich Repeat

Typically, hydrophobicity is one of the basic determinants of protein’s topology and secondary structure, as well as specific cellular localization [20]. Therefore, we performed a hydrophobic analysis of the human zyxin family that demonstrates that the LIM1 and LIM2 domains of Zyxin, TRIP6, LimD1, LPP, and Ajuba are more hydrophobic as compared to the LIM3 domain **(Supplementary Figure 1B).** In the case of FBLIM1 and WTIP, we observed that the hydrophobicity of the LIM2 domain was higher than LIM1 and LIM3 domains. The explicit compositional analysis of hydrophobic amino acids such as isoleucine, valine, leucine, cysteine, methionine, alanine and tryptophan in each Lim region imparts the variable percentage of hydrophobicity. Interestingly, only Lim3 of TRIP6 has one tryptophan which suggests that this Lim might have helix tilting **(Supplementary Table 1).** Further phylogenetic analysis of the Zyxin family proteins was carried out to understand the relatedness of the human zyxin family members. The dendrogram presented in **Supplementary Figure 2A** suggests that LimD1 is the most distant member, whereas all other members are more closely related. Furthermore, based on the evolutionary distance, zyxin appears to be the most divergent member of the Zyxin family. Amino acids sequence alignment of human zyxin protein studies suggests that zyxin family members are paralogous and could have arisen by gene duplication. In order to explore more about the gene duplication events and it’s essentiality, we performed ohnologs analysis [21]. We found that zyxin exhibits two ohnolog groups, one encompasses the FBLIM, LPP, TRIP6 and Zyxin; and another one WTIP, Ajuba and LimD1 **(Supplementary Table 2).** These results suggest that zyxin family members are infrequent to small-scale duplication (SSD) and copy number variation (CNV).

Next, we implemented NetNES [22] algorithm to determine whether these proteins feature a nuclear export signal (NES) sequence. Each protein sequence was subjected to the NetNES server, which suggested that all proteins contain NES sequences in their PRR region **(Figure 2A).** Further analysis revealed that FBLIM1/WTIP/LimD1 and TRIP6/Ajuba display LxxLxL and LxxxLxL consensus sequences, respectively. However, zyxin and LPP have some promiscuity and show the LxxxLxxL/LxxLxxL consensus sequences **(Figure 2B & 2C).** Furthermore, to investigate if the NES from human Zyxin family proteins is conserved within the vertebrates, we performed another alignment study. First, we performed BLAST analysis using the human zyxin sequence and identified 100 top scoring proteins. Subsequently, we implemented these sequences in multiple alignment functions, identified consensus NES sequences using Jalview and visualized them using WEBLOGO. This process was independently implemented for all members of the Zyxin family, which suggested that the NES sequence is conserved in each protein irrespective of their origin and function, however, the position of NES varies significantly amongst each Zyxin family member **(Figures 2A, 2B & 2C).** Since many of the Zyxin family proteins have been reported to localize into the nucleus, we also performed the nuclear localization sequence (NLS) analysis using the NLStradamus, and observed a clear presence of recognizable NLS sequence within PRR of WTIP and Ajuba **(Figure 2D).** Taken together, the results of the primary sequence analysis reveal a high extent of similarity between Lim domains of Zyxin family members. We observed variability with respect to the hydrophobicity of individual LIM domains within a given protein, as well as some variability in the position of the NES. With respect to NLS, at this point, we were able to identify its presence in two out of seven family members.

**Figure 2:**
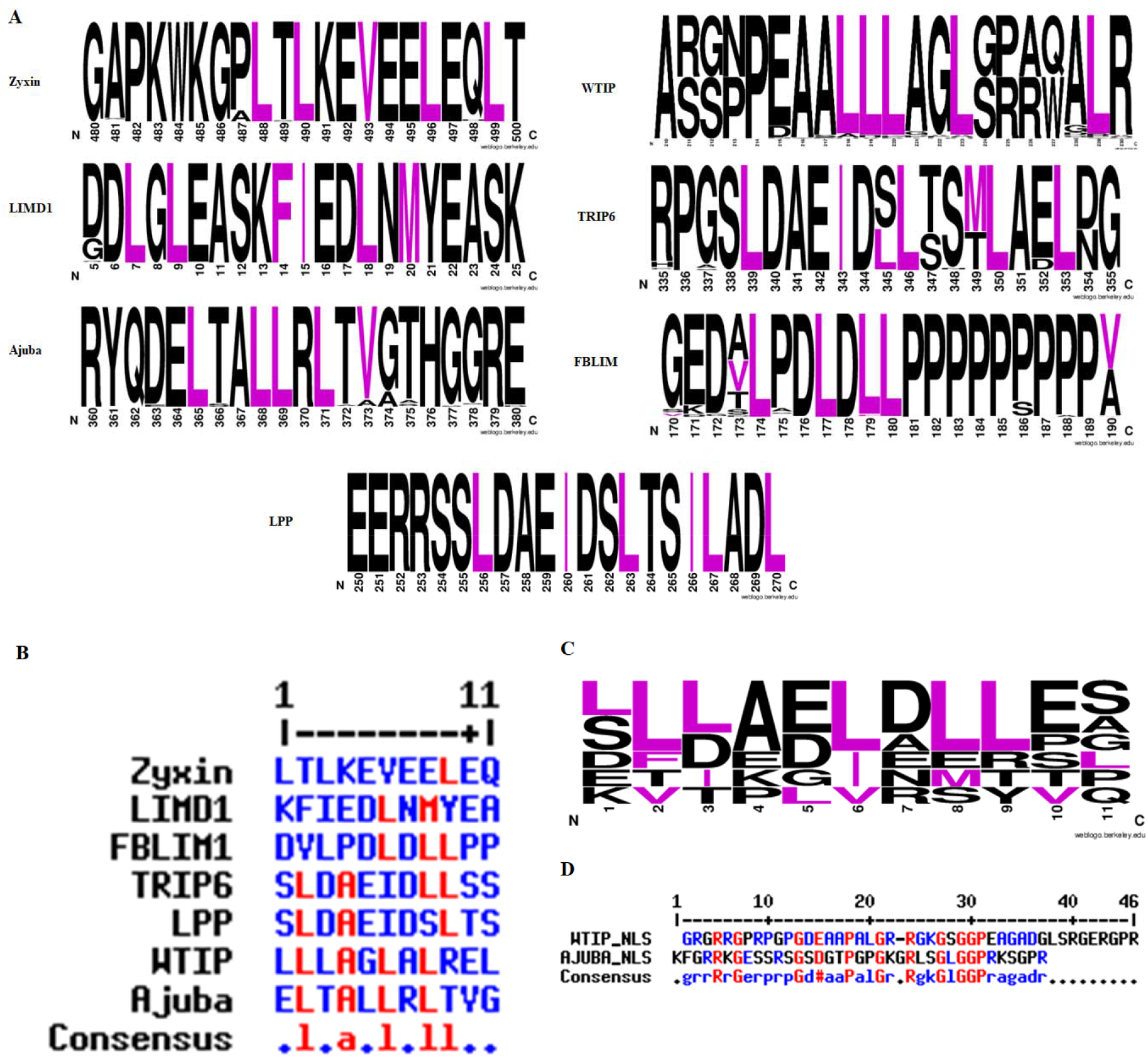
(A) The nuclear export sequence (NES) of the Zyxin family proteins. To investigate the conservation of NES in proteins from other organisms, we aligned the human protein sequence using BLAST against the non-redundant protein sequences databases. The resultant sequences obtained from BLAST were subjected to multiple alignments. (B) The NES sequence for each protein was identified through the NetNES webserver, and the sequences were aligned using MultAlin. (C) WEBLOGO was used to investigate the conservation of residues in the sequences, which suggested the presence of signature LxxLxL/LxxxLxL, (L=L/V/I/F/M; x=any amino acid) repeat, in NES sequence across all the members of Zyxin family except zyxin and LPP which shows LxxxLxxL/LxxLxxL consensus sequence. (D) The nuclear localization sequence for the zyxin family members was fetched by the NLStradamus webserver with a prediction cut-off set at a recommended value of 0.6. The conservation of predicted sequences was analysed using MulitAlin.

### 2.2. Impact of post-translational modifications (PTMs) on the isoelectric point (pI) of Zyxin family proteins

In order to understand the biophysical properties of the Zyxin family proteins and their LIM domains, we have calculated the pI of the unmodified full-length proteins as well as the LIM domain. We observed that in full-length proteins pI values range from 5.71, for FBLIM1, to 8.52, for WTIP **(Table 1**). These differences may be functionally important, as FBLIM1 is found exclusively at cell adhesion sites, and pI of 5.71 for this protein supports the fact that many acidic proteins are present in cytosol, cytoskeleton, vacuoles, and lysosomes [23]. On the other hand, WTIP is more basic, is predominantly present in the nucleus. Other Zyxin family proteins such as Zyxin, LimD1, Ajuba, LPP, WTIP, TRIP6 had calculated pI values within a single pI unit, between 6.20 and 7.19. Functionally, these proteins are found in the cell cytoskeleton and nucleus, and these pI values fall well within the range observed for other proteins with similar cellular localization [23, 24]. When this analysis was done for Lim domains only, the pI range narrowed to 5.81 to 7.81 **(Table 1).**

Next, we analyzed whether Zyxin family members display predicted PTM sites. The PTM sites we considered were acetylation, mono-methylation, ubiquitylation, and phosphorylation. This analysis indicated that phosphorylation is the predominant PTM for Zyxin family proteins **(Table 3).** Since the pI has a critical role in protein distribution inside cells and phosphorylation impacts the pI of proteins [24, 25], we also analyzed the pI of each Zyxin family proteins once phosphorylated **(Supplementary Figure 3 & Table 3).** The phosphorylation analysis of Zyxin family proteins by PhosphositePlus [26] indicated that the pI shift varied depending on the number of phosphorylation modifications and affects the pI considerably **(Table 3).** For example, the pI of zyxin without PTMs is 6.22, while calculated pI values ranged from 3.17 to 6.12, depending on the type of PTM. The most dramatic change was observed for LPP, a Zyxin family member mainly involved in cell motility and gene transcription [27]. LPP was predicted to undergo the drastic range of a pI from 7.18 for unphosphorylated to 2.84 for a completely phosphorylated state with all 83 proposed phosphorylation sites. Interestingly, the FBLIM1, the most acidic of Zyxin family member in non-PTM form, undergoes the least amount of phosphorylation, hence, its pI is not affected drastically **(Table 3).**

**Table 3:**
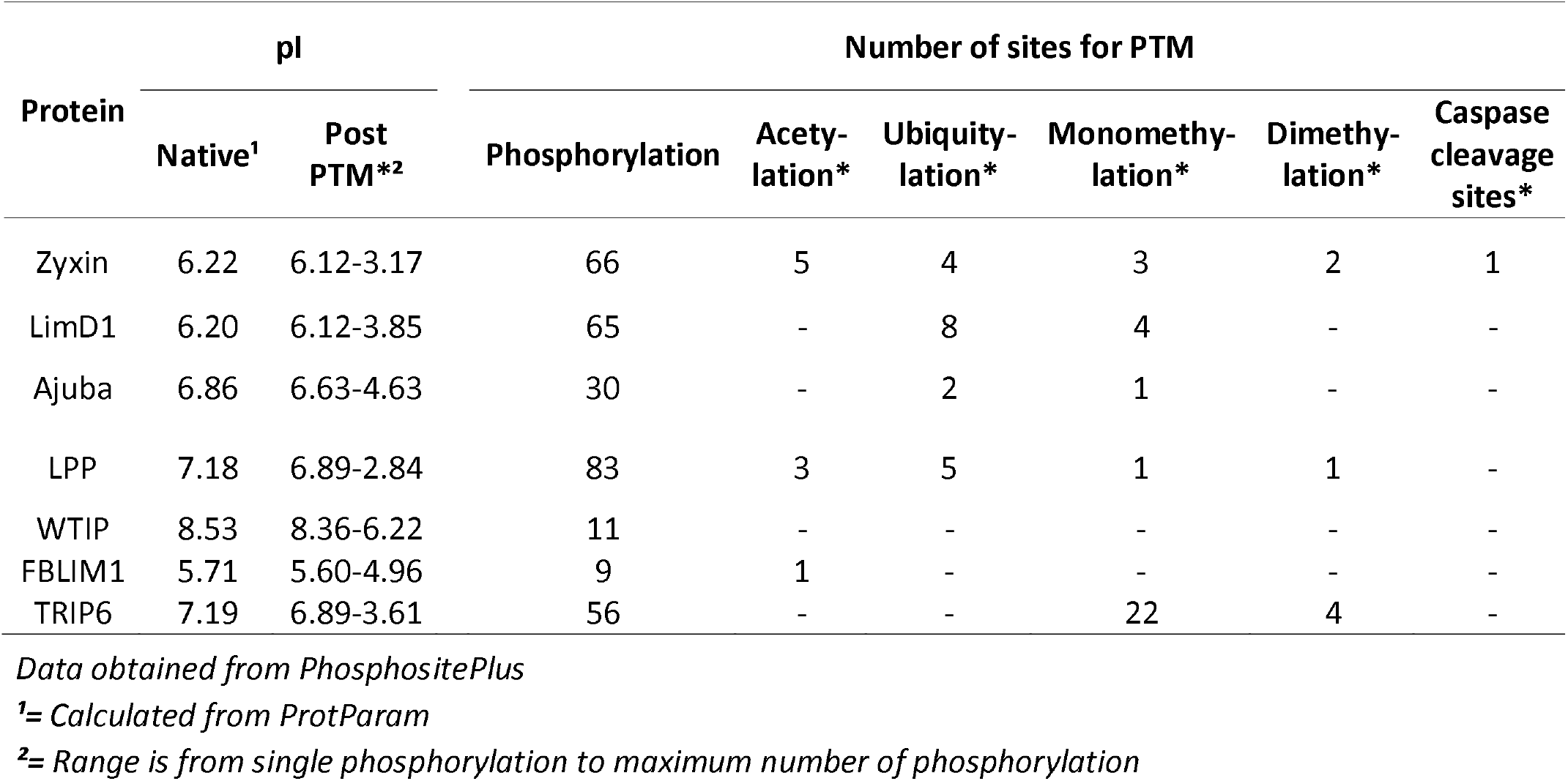
Effect of post-translational modifications on isoelectric point of Zyxin family proteins.

Overall, this analysis suggests that Zyxin family members display differences in their pI values, which are likely linked to their function. Additionally, we predict that all these proteins may be regulated by different PTMs. The primary PTM of the Zyxin family might be phosphorylation, and we observe a significant impact on pI values as a consequence of this PTM.

### 2.3. Structural features of Zyxin family Lim domains indicate their similarities with transcription factors

After exploring the sequence of Zyxin family proteins, we further sought to investigate their structural similarities and dissimilarities. Due to the absence of high-resolution structures of Zyxin family Lim domains, we decided to perform homology modeling for each LIM domain. The modeling was performed using Robetta package and models were generated for all three LIM domains linked together, not individually (see Methods for details). The homology modeling suggested that LIM1-3 adopt a hairpin shape, placing N- and C-terminus into proximity in all members of the family, except for Ajuba and TRIP6 **(Figure 3).** In the case of TRIP6, both the terminals were positioned diametrically opposite to each other, whereas for Ajuba, the C-terminus is located in the middle, and the N-terminus is present at the opposite end **(Figure 3).** Model validation using PROCHECK analysis, which employs the Ramachandran plot, suggested that all the residues were in permitted regions and display acceptable stereochemistry **(Supplementary Figure 4A)** [28]. Absolute quality estimation is another parameter that allows a comparison of the modeled structure with reported PDB structures. The score of comparison is represented via Z-score, where a Z-score in between 0.5 to 1 represents a model that closely resembles an experimentally determined native structure **(Supplementary Figure 4B**)[29]. Quantitative Model Energy ANalysis (QMEAN) of LIM domain models of Zyxin family proteins showed a Z-score ranging from 0.6 – 0.7, which is well in the acceptable range of QMEAN Z-score. Thus, the comprehensive model validation performed using PROCHECK and QMEAN analysis showed that all models have a good quality of stereochemistry as well as nativeness, respectively, making them reliable for further studies.

**Figure 3:**
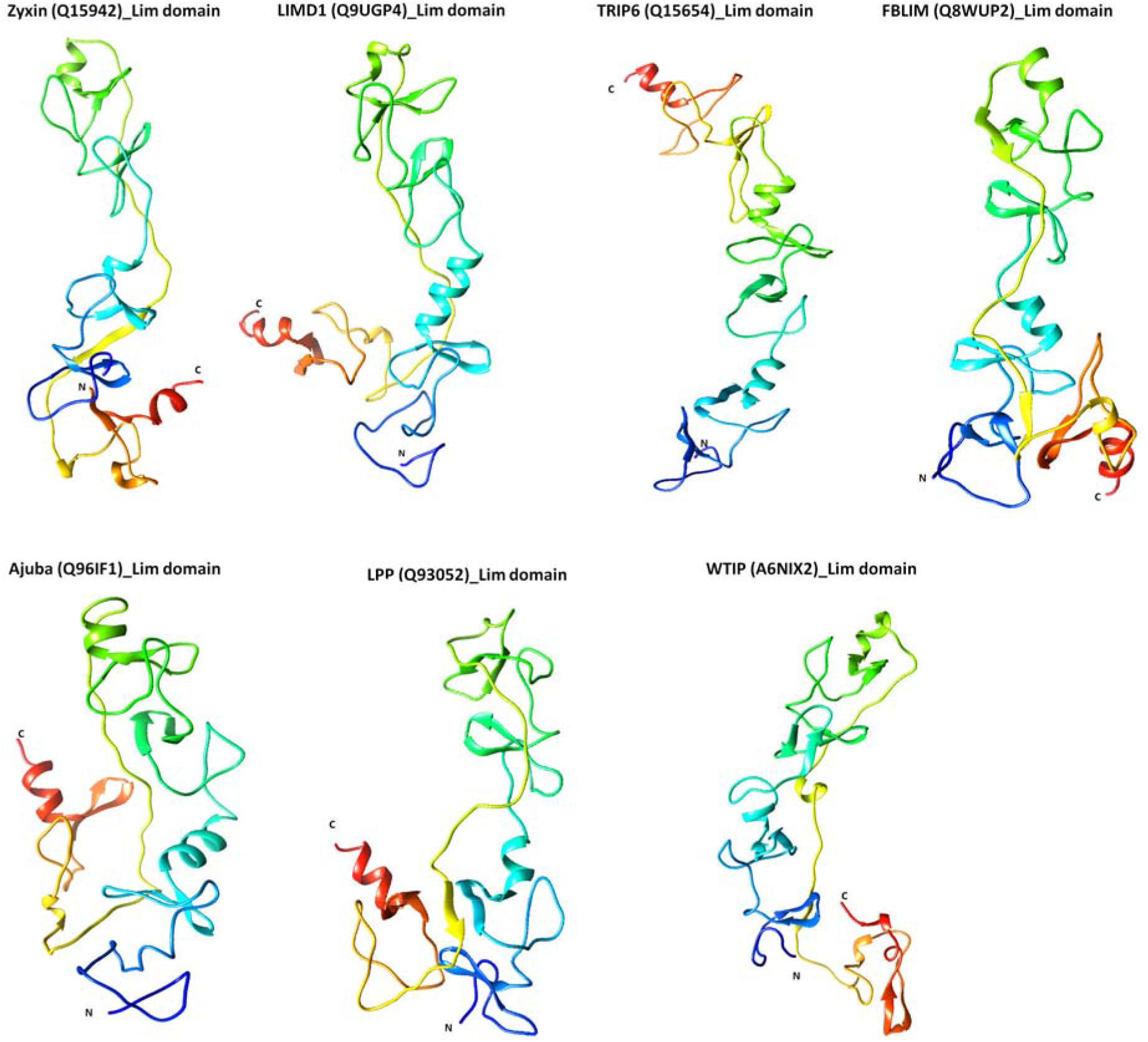
(A) Structures of the Lim domains of the Zyxin family proteins. The structures were generated using comparative/homology modeling in the Robetta server. Robetta server generates structure using structures identified by BLAST and PSI-BLAST, the regions of the protein that could not be modeled using BLAST are modeled de-novo using Rosetta fragment insertion method. For each model, the N- and C-terminal are depicted in blue and red, respectively. All models were validated using PROCHECK and SWISS-MODEL workspace; and the structures with the highest Z-score are presented in Supplementary Figure 4.

As LIM domains of zyxin and TRIP6 have been shown to interact with nucleic acids [30] we utilized DNAbinder webserver [31] to investigate if other Zyxin family proteins can bind to nucleic acids. DNAbinder trains a Support Vector Machine (SVM) using a database to search for sequence similarity and motif finding approaches to discriminate DNA binding proteins from non-binders. Our analysis indicated that all the members of the Zyxin family have positive SVM scores ranging from 0.11 (LPP) to 1.71 (WTIP), suggesting that all the proteins can potentially interact with DNA **(Table 4).** Furthermore, we used DALI server [32] to investigate whether Zyxin family proteins are structurally similar to known DNA binding proteins. Our alignment of the homology models of LIM-domains of each Zyxin family protein **(Table 5)** with known structures from DALI server led to the identification of ~88 structures. Interestingly, the majority of the structures shared only ~15 – 30 % sequence identity (% id), however, the Z-score values demonstrated high structural similarities with established transcription factors presented in **Supplementary Figure 5.** The Z-score depicts the similarity between the protein of interest with the proteins from the PDB and the Structural Classification of Proteins (SCOP) databases. Ultimately, the inclusion of both databases helped us to remove biases introduced by size, substructures and divergence, which may result in the server giving false-positive proteins showing similarity [32, 33]. Thus, we decided to further study the top 6 structures with a Z-score >9.4, while exhibiting only 23–28 % sequence identity **(Supplementary Figure 5**). The highest scoring structure in DALI analysis was a complex of Lmo4 protein and Lim domain (PDB ID: 1RUT) [34], followed by a complex of Lim Only protein 4 (LMO4) and Lim domain-binding protein (LDB1) (PDB ID: 2DFY) [35], the complex of Lim domain from homeobox protein LHX3 and islet-1 (ISL1) (PDB ID: 2RGT) [36], Lim/homeobox protein Lhx4, insulin gene enhancer (ISL2) (PDB ID: 6CME) [37] and an intramolecular as well as the intermolecular complex of Lim domains from homeobox protein LHX3 and islet-1 (ISL1) (PDB ID: 4JCJ) [38] (see **Supplementary Figure 5 & Table 5).** The highest scorer 1RUT which included Lim domain transcription factor LMO4 (Lim Only) was first identified as a breast cancer autoantigen [39], which works as a transcriptional factor [40] and interacts with tumor suppressor Breast Cancer protein 1 (BRCA1) [41]. The expression of LMO4 is developmentally regulated in the mammary gland and it is known to repress the transcriptional activity of BRCA1 [41]. A common theme observed in all the highest scores protein complexes was that the participating Lim domains were part of transcription factors critical for developmental pathways. The structural similarities of Zyxin Lim domains with various transcriptional regulators suggest it may have a role in nucleic acid binding and its regulation.

**Table 4:** SVM based prediction of DNA binding of Zyxin family proteins as calculated by DNAbinder webserver

| Protein | SVM score* | Prediction | Job Number on webserver |
| --- | --- | --- | --- |
| Zyxin | 1.2919473 | DNA binding protein | 1710 |
| LimD1 | 0.90588128 | DNA binding protein | 4069 |
| Ajuba | 1.4081202 | DNA binding protein | 7780 |
| LPP | 0.11482763 | DNA binding protein | 1668 |
| WTIP | 1.7520286 | DNA binding protein | 1032 |
| FBLIM1 | 0.39872591 | DNA binding protein | 1215 |
| TRIP6 | 0.83398047 | DNA binding protein | 3994 |

*\*= Calculated for full length (all amino acids) under amino acid composition mode*

**Table 5:**
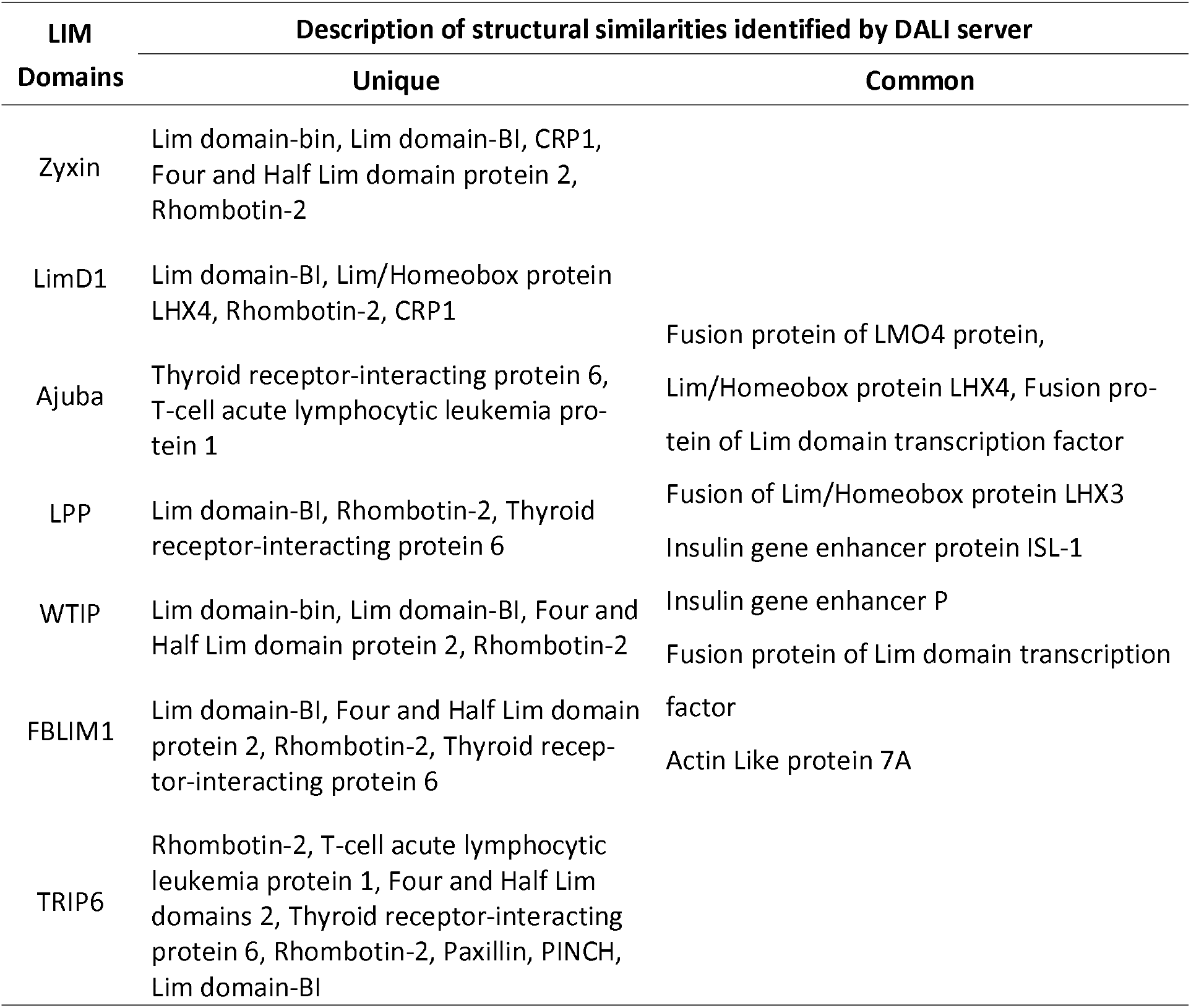
DALI structural alignment of the LIM domain of Zyxin proteins

To further explore the possibility that Zyxin family members Lim domains bind nucleic acids directly, we used our homologous models and mapped electrostatic surface potential using Adaptive Poisson-Boltzmann Solver (APBS) and Yellow-Red-Blue (YRB) analysis as a strategy to explore whether these structures have positively charged regions and nucleotide hydrophobic regions, respectively **(Supplementary Figure 6).** It showed that zyxin, Ajuba, LPP and WTIP have higher positive electrostatic patches with hydrophobic regions which suggest that these might be contributing to the nucleic acid-binding.

### 2.4. Zyxin family Lim domains display contrasting dynamics with large inter-domain motion of individual Lim1/2/3 domains

As our homology models indicate, individual Lim domains do not share extensive similarities making it difficult to rationalize the diverse range of functions they exhibit. In addition to structure, the intrinsic dynamics of proteins is an important determinant of function. Therefore, to understand the intrinsic dynamics of Zyxin family LIM domains, we performed NMA (**N**ormal **M**ode **A**nalysis) and Covariance analysis of LIM domains using iMODS [42]. NMA of Zyxin family LIM domains suggested that all proteins show a pinching motion **(Figure 4).** In order to make a finer observation of the movement, we performed Covariance analysis which indicates the variability of correlated, uncorrelated, and anti-correlated motions. The covariance matrix provides information on the fluctuations for C-alpha atoms around their mean positions and coupling between pairs of the residues [43]. **Figure 5A** presents correlated (red region), uncorrelated (white) and anti-correlated (blue) motions for the Zyxin family LIM domains, which suggests that LIM1 had the most correlated motion across all the members. In contrast, LIM2 domain of all the members displayed varied motion **(Figure 5A).** We also noticed that LIM3 of Ajuba, LPP and WTIP have similar correlated motions. The Covariance matrix also indicated that TRIP6 has the highest correlated motion whereas Ajuba and WTIP have relatively lower correlated motion. Furthermore, we also observed that Ajuba and WTIP, as well as LPP, LIMD1, and FBLIM have similar motion but Zyxin and TRIP6 have a completely unique motion amongst the Zyxin family proteins **(Figure 5A).** Next, we employed iMODs for Elastic Network Model (ENM) analysis of Zyxin family LIM domains to identify the pairs of atoms that are connected by springs. The results are depicted by a grayscale graph, where each dot represents one spring between two atom pairs. The degree of greyness for each dot represents the stiffness, dark grey represents stiffer springs and *vice versa*. LIM1 domains of all the Zyxin family proteins presented higher stiffness as compared to the LIM2 and LIM3 regions **(Figure 5B).** Note that the stiff grey regions of LIM1 domains also present correlated motions **(Figures 5A & 5B).** Overall, we observed that LIM1-3 in all Zyxin family members display “clamping” motion in our NMA suggesting that these large-scale motions may be functionally relevant. However, covariance analysis and ENM analysis revealed variability among the family members, which is broadly in agreement with our expectation that the functional differences may be due to a difference in intrinsic dynamics.

**Figure 4:**
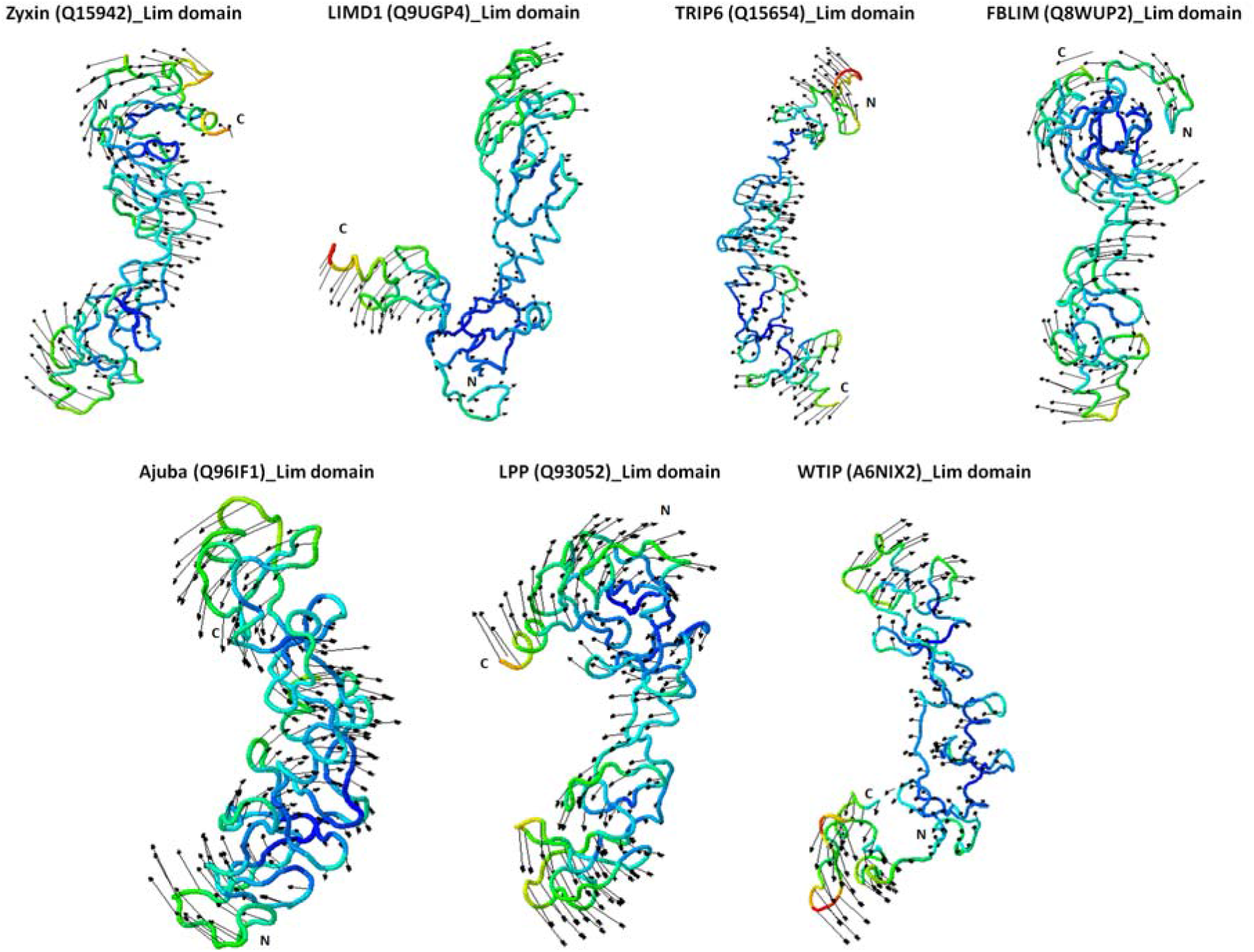
Normal mode analysis (NMA) of Zyxin family Lim domain proteins. The NMA for each protein model was performed using the iMODS server under default settings. The motion of each domain is indicated by the arrows arising in the direction of motion. Additionally, the models are coloured based on NMA mobility of each structure, where the mobility is represented using a colour spectrum (blue represents the lowest mobility whereas red represents the highest mobility, and intermediate mobility is indicated by interim colours, blue>green>yellow>red). Interestingly, the direction of motion of all proteins is such that the distal ends of the protein move in the direction towards each other producing a motion that resembles clamping.

**Figure 5:**
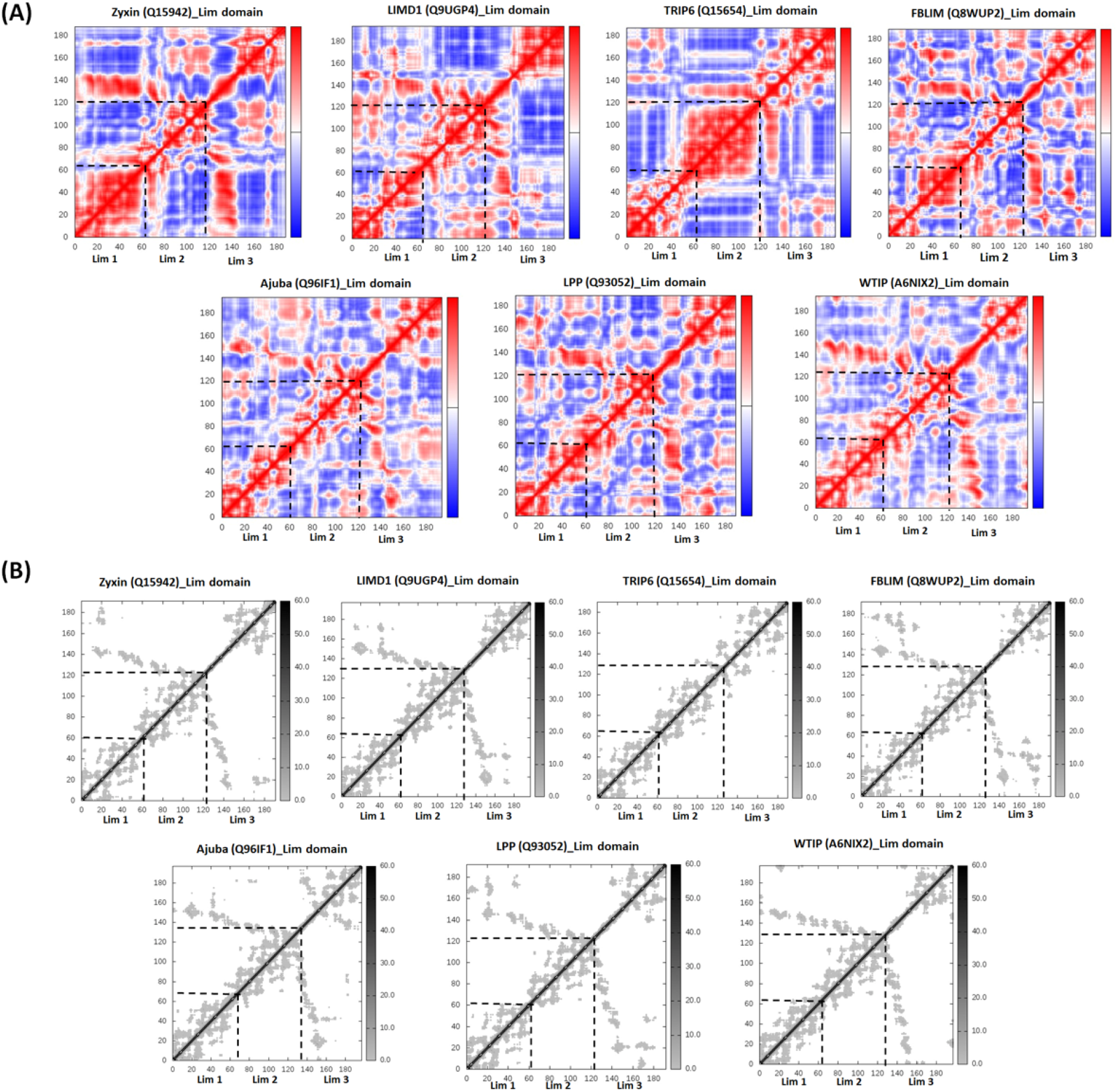
(A) Covariance analysis of the Zyxin family Lim domain. Covariance analysis was performed to understand the correlated movement of protein residues in the secondary structure which can shed light on the function of the protein. Each panel represents the covariance matrix for each member of the Zyxin family, in the matrix correlated motion is depicted as red, anti-correlated as blue, and uncorrelated as a white dot. High levels of correlated motions were observed in case of Lim1 domains of all protein proteins, varied amount of motion was observed in case of Lim2 with the highest in TRIP6, again Lim3 showed a similar kind of motion in all proteins. Proteins with similar localization and function demonstrated similarity in motion. For example, Ajuba, LIMD1, and WTIP displayed relatively similar covariance matrix of Lim3 domain these proteins are localized at the same site and constitute P-Body components, primarily perform gene regulation and mediate miRNA silencing. LPP and FBLIM1 showed similar correlated, uncorrelated and anticorrelated motions in Lim1/2/3, and both proteins involved mainly in cell shape maintenance; supporting the fact that similar dynamics may have similar localization and functions, each Lim region has its own characteristic features and believed to perform functions independently. (B) The linking matrix of the Elastic network model demonstrates the stiffness of the springs connecting the corresponding pair of atoms. The dots are coloured as a function of their stiffness, darker colours reflect stiffer springs. Interestingly all proteins showed a similar kind of stiffness except TRIP6.

### 2.5. Mapping of protein-protein interactions of the Zyxin family proteins provides insights into their crucial role in cancer progression

The Zyxin family proteins are functionally diverse, and many of their functions are attributed to their abilities to interact with a wide range of host proteins. Therefore, we decided to map the protein interactome for the Zyxin protein family members using databases such as Genevestigator®, PhosphoSitePlus®, Bioplex [44], GeneMANIA [45], MINT [46], STITCH [47], Signor 2.0 [48] and STRING [49]. First, we used the Genevestigator database that provides information on the protein expression levels in various cancers. As presented in **Supplementary Figure 7 & 8,** where the x-axis represents the protein expression level, the Zyxin family proteins are upregulated in almost all cancer types, including breast, colon, kidney, liver, and prostate cancers. We next performed a mutation frequency analysis using PhosphoSitePlus databases, which allowed us to examine the effect of mutated protein on cellular functions, pathologies and diseases [50]. The resultant lollipop plots for each Zyxin family proteins are presented as a function of particular cancer types in **Figure 6**. These plots suggest Zyxin, LPP, LIMD1, TRIP6, and FBLIM display high levels of mutational frequency in stomach cancer **(Figure 6).** Similarly, Zyxin, LIMD1 and TRIP6 mutations are linked with endometrial cancer. In the case of lung cancer, LPP displayed the highest amount of mutation frequency. For bladder, kidney and head/neck cancers, proteins TRIP6, WTIP, and Ajuba, respectively, showed the highest mutational frequency.

**Figure 6:**
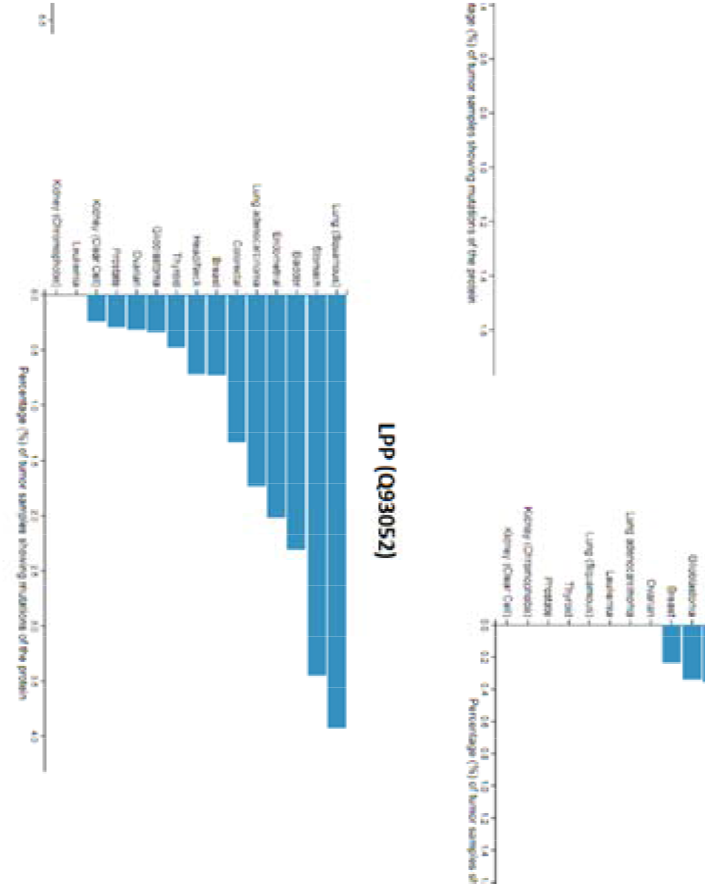
Mutational frequency profiles of the Zyxin family proteins with different cancer malignancies. All proteins exhibit a considerable amount of mutational frequency which varied for each member. The highest mutation frequency was observed for Ajuba and the lowest was observed for LIMD1.

Using Bioplex, we identified different cytoskeleton and gene regulatory proteins that interact with Zyxin family proteins. Bioplex depicts the protein-protein interactions and also the probabilities of association using the HEK293T cell line as a model. **Supplementary Figure 9–14** presents the interacting proteins as prey (green circle) and bait (grey square) and the direction of interaction is represented by the arrow. Interestingly, we observed that the Zyxin family proteins interact amongst themselves as well, with Ajuba being the focal point showing interactions with TRIP6, LIMD1, Zyxin, and LPP. WTIP and FBLIM1, however, do not indicate such interactions. Also, zyxin interacts with VASP, FHL3, and TANC2 **(Supplementary Figure 9–14).** LimD1 and TRIP6 act more closely to regulate the interaction network amongst themselves. TRIP6 has the most complex interactome compared to other Zyxin family members **(Supplementary Figure 14).** Alternatively, WTIP displayed the smallest interaction network consisting of two proteins only (TRIP6 and PPP2R3A).

Next, we performed physical association, co-expression, and pathway analysis for the Zyxin family proteins using GeneMANIA, which manifested the involvement of each Zyxin family protein as a critical player that could alter the signaling pathways **(Figure 7).** Zyxin, LimD1, and Ajuba demonstrated more physical interactions as compared to WTIP, LPP, and FBLIM1. Moreover, zyxin, FBLIM1, Ajuba, and TRIP6 indicated higher degrees of co-localization/co-expression with other proteins. LPP demonstrated a direct and indirect co-expression with many genes, particularly TAGLN, CNN1, and SMTN **(Figures 7).** Using this pathway analysis, we found that zyxin could be linked with NOLC1 (Nucleolar and coiled-body phosphoprotein 1), which facilitates ribosomal processing and modifications [51]. Pathway analysis also suggests that zyxin, FBLIM and LPP interact with VASP, indicating their similar roles or compensatory nature where the absence of one of the proteins can be compensated by another protein of the same family. Pathway analysis by OmniPath identified the role of Zyxin in the AKT and IL7R pathways **(Supplementary Figure 15),** which was further supported by the Signor analysis, which found that Akt phosphorylates Zyxin on Ser142 and regulates apoptosis [52].

**Figure 7:**
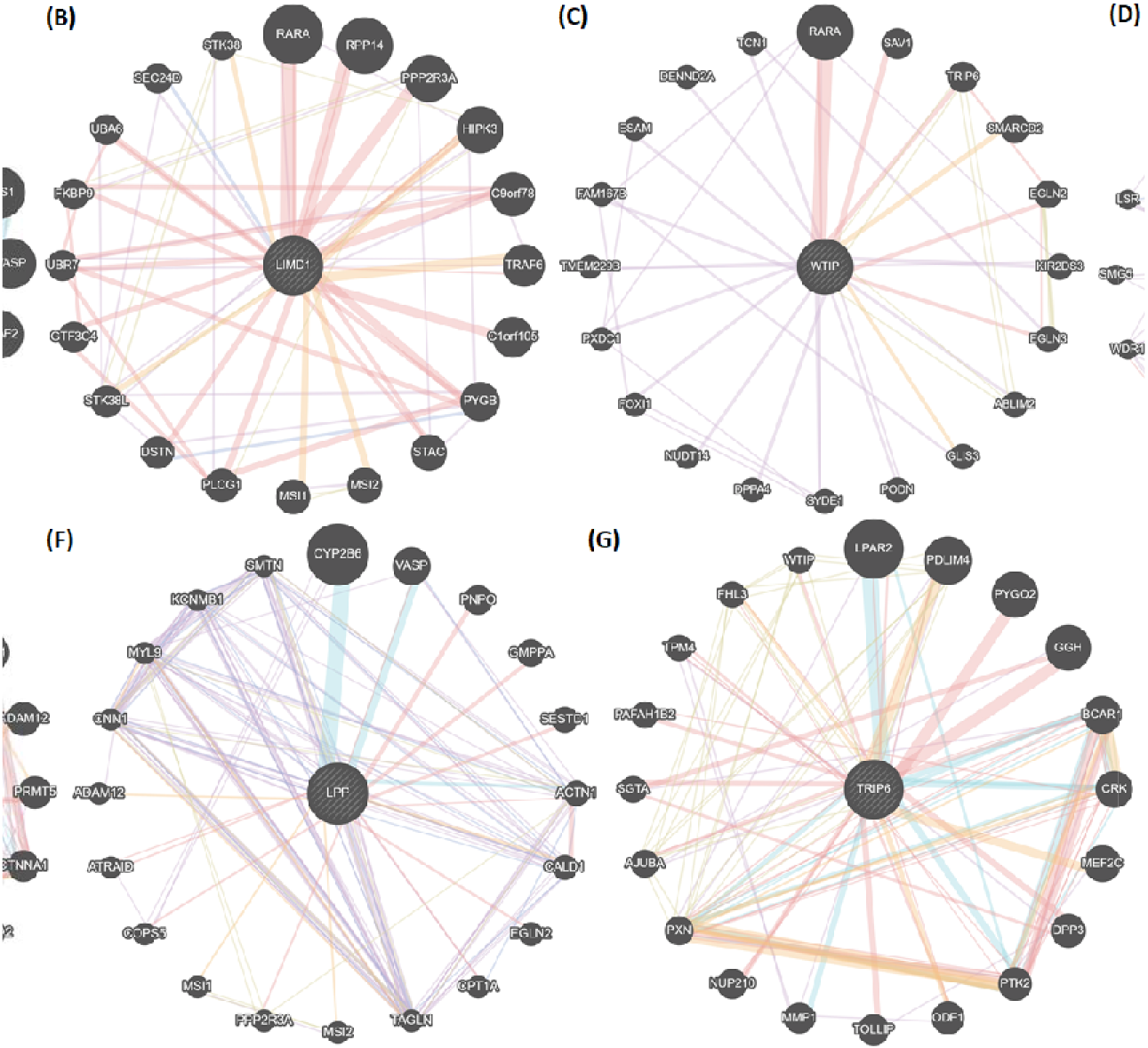
Pathway analysis by GeneMANIA. GeneMANIA highlighted the pathways that each protein member interacts within a cell. This helped us identify various proteins that interact with the query protein. The connections are shown by blue lines, the width of each line represents the degree of interaction through weighted representation. The thicker line represents more interaction and vice versa.

We have also used STITCH (Search Tool for Interactions of Chemicals), STRING (Search Tool for Retrieval of Interacting Genes/Proteins) and MINT (Molecular INTeraction database) databases to mine the protein-protein interactions of the Zyxin family proteins **(Supplementary Figure 15 & Table 6).** STITCH database provides information on interacting partners of a protein of interest, which could, in turn, provide a further understanding of their molecular/cellular functions [47]. On the other hand, the STRING database provides enhanced coverage of protein associations with functional genome-wide discoveries [49]. Upon performing the STITCH/STRING analysis, we observed that zyxin interacts with BCAR1 (Breast cancer anti-estrogen resistance protein 1), which coordinates tyrosine kinase-based signaling [53, 54].

**Table 6:**
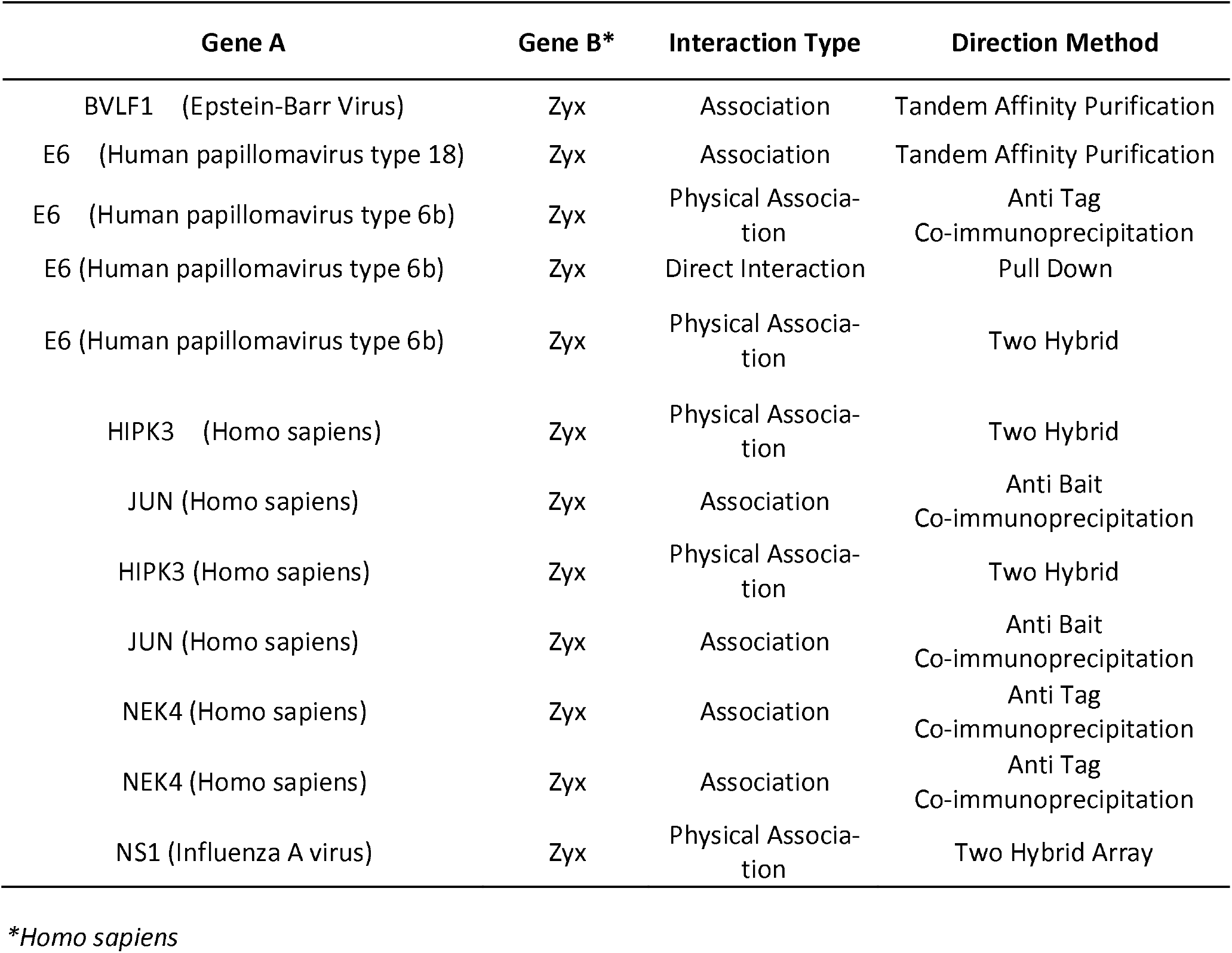
MINT analysis of Zyxin protein

## 3. Discussion

In this work, we report the results of a comprehensive bioinformatic analysis of the human Zyxin family proteins. We performed primary sequence analysis, homology modeling, analysis of protein dynamics and interactions. The multiple sequence alignment of LIM domains suggested that although key Zn^2+^ ion coordinating ligands and the spacing between them seems to be relatively conserved, the overall sequence conservation, including the hydrophobic residues is relatively low **(Supplementary Figure 1).** This is in agreement with subsequent hydrophobic analysis, which revealed that the LIM domains have different hydrophobicity levels **(Figure 1C).** The explicit compositional analysis of hydrophobic amino acids indicated the variable percentage of hydrophobicity **(Supplementary Table 1).** The presence of tryptophan in the only Lim3 of TRIP6 has intrigued further to Lim domain functions, because tryptophan plays an important role in hydrophobic-hydrophilic interfaces, hydrophobic mismatch and helix tilting, and has the importance in the binding of other non-proteinaceous biomolecules [55, 56]. Such differential hydrophobicity in LIM domains could be linked to their distinct properties, structures, and functions, despite them sharing conserved zinc-finger organization. Furthermore, the calculated pI values suggested the presence of two main clusters of Zyxin family proteins, those with pI lower than 7.5 (Zyxin, TRIP6, LimD1, and LPP) and those with pI value higher than 7.5 (FBLIM1 and WTIP). Overall, these findings are consistent with the bimodal distribution of acidic and basic proteins which in turn is correlated with the subcellular localization and cellular niches [23].

PTMs are major regulators of protein function and, in many cases, the presence of PTMs changes the properties of the target protein. Our analysis indicated the presence of many phosphorylation sites, which are located in the PRR of all the proteins **(Supplementary Figure 3 & Table 3)** and the influence of the extent of phosphorylation on the pI. Therefore, these results suggest that phosphorylation, and the changes in pI it generates, may be linked with protein cellular localization and functional differences, as observed in other systems (Alende, Nielsen et al. 2011).

Zyxin family members appear to be paralogous and might have arisen by gene duplication events. Ohnolog analysis indicated that zyxin family proteins are retained during 2-rounds of whole-genome duplication events (2-WGD) which suggests its essentiality to vertebrates. Ohnologs were about 30% protein-coding genes that are retained from two rounds of genome duplication (2R-WGD) events [57]. Ohnologs were known to be involved in gene enrichment and dosage compensation, and it is retained in vertebrates due to its essentiality and associated with dosage imbalance diseases such as down syndrome and cancer [21, 57]. Another feature that could have a major impact on the intracellular localization of the Zyxin family members is the presence of the NES and/or NLS sequence. We demonstrate that the consensus sequences are present in all of the Zyxin family proteins irrespective of their reported function and localization **(Figure 2A, 2B & 2C).** The NES consensus sequence we identified and their frequency of occurrence are in good agreement with the available report that suggests that out of all the NES containing proteins, 75% have LxxLxL, 11% have LxxxLxL and 15% of proteins do not comply with either of these consensus sequences [22]. We also analyzed the primary sequences of the Zyxin family for the presence of an NLS **(Figure 2D).** Here, our analysis identified a consensus NLS in only two (out of seven) proteins analyzed suggesting that further studies are needed to better understand Zyxin family members cellular distribution (localization) at a more molecular level and mechanistic level. We would also like to stress that our phylogenetic and NES analysis supports the hypothesis that the Zyxin family proteins are evolved from a common ancestor and have retained the NES sequence, and diversification in these proteins resulted in the emergence/shift of functions from the nuclear roles to cytoskeletal properties [58].

Previous work has suggested that Lim domains can serve as a protein-binding interface and also interact with nucleic acids [17–19, 30, 59–63]. Our homology modeling yielded structural models of sufficient quality to enable additional analysis, including broad structural alignment that identified transcription factors as most closely structural related proteins to Zyxin family members **(Figure 3, Supplementary Figure 4A, 4B & 5).** These results prompted us to hypothesize that Zyxin family members could bind nucleic acids, primarily DNA. We probed this question further by making the electrostatic surface potential onto homology models, which revealed that 5 out 7 models contain a positively charged surface patch suggestive of nucleic acid binding capacity **(Supplementary Figure 6).**

An additional observation we made upon inspecting our homology models is that Zyxin family of proteins LIM1-3 domains are relatively structurally similar leading us to examine whether well-documented functional differences are mostly due to differences in intrinsic dynamic, rather than structure. Our analysis of correlated motions, which are important for processes like allosteric regulation, catalysis, ligand binding, and protein folding [64–67], revealed that LIM1 displayed correlated motion, therefore suggesting that LIM1 domains will exhibit a higher propensity for binding towards its binding partner **(Figure 5).** However, the Lim3 region of each protein shows more anti-correlated motion. These transformations in the covariance matrix of Lim domains reflect the differences in the internal structural dynamics of the Zyxin family proteins, and these dynamic behaviours could confer diverse functions to each Zyxin family member as well as each LIM regions. Similarly, stiffness of the protein was investigated using ENM. The observations made for stiffness were in corroboration with the covariance analysis: the stretches that showed the highest correlated motion also showed a high degree of stiffness **(Figure 5A & 5B).**

Lastly, we observed that Zyxin family proteins are upregulated and have a high mutational frequency in a variety of cancers **(Figures 6 & Supplementary 7, 8),** in agreement with the large amount of literature that suggests their involvement. Our protein-protein interaction (PPI) analysis generated a closer view of the PPIs that these proteins are involved in. The results of PPI analysis show that each family member establishes a different (function-dependent) network of interaction partners. In some cases, individual interaction networks include other zyxin family members, and feature a diverse range of partners. On the other hand, the WTIP PPI network was limited to only two interaction partners, including a Zyxin family protein TRIP6. These PPI networks could provide new insights into the functional diversity of the Zyxin protein family. For example, the interaction of Zyxin with NOLC1 (Nucleolar and coiled-body phosphoprotein 1) and BCAR1 (Breast cancer anti-estrogen resistance protein 1) suggest a novel role of zyxin in ribosomal processing and in tyrosine kinase-based signaling, respectively **(Figures 7, Supplementary Figures 9–15).** Additionally, we observed that many of the interacting partners play a role in the rewiring of the cancer metabolism and contributing to oncogene addiction, which has important implications for understanding the role these proteins play in cancer. Going forward, these PPI networks are likely to reveal additional relationships and connections that justify future follow up studies.

From a broader perspective, our bioinformatic study yields new insights into an incompletely understood family of LIM-domain containing proteins, the Zyxin family. Our results clearly indicate that these proteins can function in a variety of ways from mediating protein-protein interaction to potentially serving as nucleic acid-binding platforms. Additional work will be required to fully investigate these emerging aspects of this family of proteins. Given the demonstrated link to human disease, we expect that the current and any future efforts will open opportunities for therapeutic development.

## 4. Materials and Methods

### 4.1 Sequence, amino acid composition, phylogeny, and basic biochemical analysis

All the human Zyxin family (Zyxin, LimD1, Ajuba, LPP, WTIP, FBLIM1, TRIP6) sequences were retrieved from the UniProt database as outlined in **Table 1** [68]. Amino acid sequence alignments were performed using MUltiple Sequence Comparison by Log-Expectation (MUSCLE) which also offers identification of critical amino acid residues [69]. Sequence PSI-BLAST (Position-Specific Iterative Basic Local Alignment Search Tool) was performed to find distant evolutionary related proteins. For the phylogenetic analysis, Zyxin family sequences were subjected to Clustal W [70], the output was then imported into Jalview [71] and a tree was generated through neighbour joining using a BLOSUM62 [72]. Ohnologs analysis was performed using OHNOLOGS v2. It is a comprehensive database for the genes which are retained from 2R-WGD and 3R-WGD. Ohnologs families for *Homo sapiens* from 2R WGD and the criterion was used for q-score for outgroup <0.01 OR q-score for self synteny <0.01. NES prediction analysis was performed using NetNES [22] which calculates the scores from the Hidden Markov Model and Artificial Neural Network. To determine the consensus sequence, conserved Leucine-rich NES signal analyzed from NetNES [22] aligned by MUSCLE [73] and logo was generated using WEBLOGO [74]. NetNES of each member of the Zyxin family was performed using pBLAST and aligned file were subjected to WEBLOGO [74]. NLStradamus webserver was used for Nuclear Localization Sequence (NLS) search [75]. The sequence of each protein was imported on the NLStradamus and the NLS prediction was run on default parameters with prediction cut-off set at a recommended value of 0.6. The conservation of predicted sequences was analysed using MultAlin [76]. ProtParam was used to calculate amino acid composition and basic biochemical characteristics [77]. We used the program ProtScale to perform hydrophobicity analysis [77]. Furthermore, PhosphoSitePlus v6.5.9.1 was used to calculate the number of phosphorylation modification sites as well as to determine the isoelectric point in the different phosphorylated states [26].

### 4.2. Structure modeling, validation and alignment of Zyxin family Lim domains

Zyxin family Lim domain amino acid sequence was retrieved from UniProtKB [68], as UniProt IDs detailed in **Table 1** were utilized to build homology models using the Rosetta server [78]. The Robetta server performs the comparative modeling with the available structure using BLAST, PSI-BLAST, and 3D-Jury packages or employs *de novo* modeling approaches using the *de novo* Rosetta fragment insertion method [79]. Next, all the Lim domain models of Zyxin family proteins were validated by SWISS-MODEL workspace [80, 81] encompassing the Anolea, DFire, QMEAN, Gromos, DSSP, Promotif, and ProCheck packages. Structure alignment of Zyxin family Lim domains was performed by the Dali server [32, 82]. DNAbinders was used to identify potential DNA binding proteins [31]. In this, we implemented the analysis using amino acid composition mode, the webserver performed a PSI-BLAST analysis against a database containing DNA binding and non-binding proteins (1153 for each case). The SVM scoring was performed against a threshold of 0, with the resultant SVM indicating if the protein interacts with DNA or not. The negative SVM values suggest non-DNA binders whereas proteins with positive SVM scores are DNA binders. Adaptive Poisson-Boltzmann Solver (APBS) electrostatics surface potential calculations were performed using APBS plugin in Pymol [83].

#### Normal mode analysis (NMA), Elastic network modeling and Covariance analysis

NMA and Covariance analyses were performed using the iMODS server using default settings [42]. After structure submission in PDB format or PDB ID, iMODS calculates the lowest frequency modes (normal mode represents the biological motions) in internal coordinates of a single/trajectories structure. This server calculates the lowest frequency normal modes in internal coordinates and utilizes the improved faster NMA calculations by implementing the iterative Krylov-subspace method [84]. iMODS also provides motion representations with a vector field, deformation analysis, eigenvalues, and covariance maps. It also has an affine model-based arrow representation of the dynamic regions [42]. Furthermore, the iMODS server also helps us understand how correlated the motion of residues in the protein is as well as how flexible the movement of each residue is under the NMA.

### 4.3 Protein-protein interactions of Zyxin family proteins

To explore the interactome of the Zyxin family proteins, we have used Bioplex explorer (Bioplex 3.0) [44]. Protein-protein interactions define the involvement of the desired protein in a particular pathway that would help to map the pathogenic network. We have utilized the Pharos database to extract the role of Lim domains in cancer and autoimmune diseases [85]. Pharos interface provides the collective information of the query protein by extracting the information from other databases, particularly in a disease context. Pharos is helpful in browsing the relevant information of the desired target in a systematic manner [85]. To get the information about the signaling pathways influenced by the Zyxin family proteins, we have used OmniPath [86] and Signor 2.0 [48] programs. We also used Genemania [45] that provides insights into protein-protein, protein-DNA and genetic interactions, pathways, gene and protein expression data, protein domains, and phenotypic screening profiles. We also explored the experimentally determined and predicted protein-protein interactions using the STRING database [49]. Next, we performed an analysis of the chemical and protein interactions using the program STITCH [47]. We also utilized the **M**olecular **INT**eraction database (MINT) database to identify the interactions mediated by the Zyxin family members [46]. MINT focuses on experimentally determined protein-protein interactions mined from the available literature. Next, we used GENEVESTIGATOR to analyze transcriptomic expression data from repositories [87]. The collection of gene expression data from different samples (tissue, disease, treatment, or genetic background) with graphics and visualization tools [87].

*For the summary of methods and servers, please refer to supplementary Table 3*.

## Supplementary Materials

**Supplementary Figure 1:**
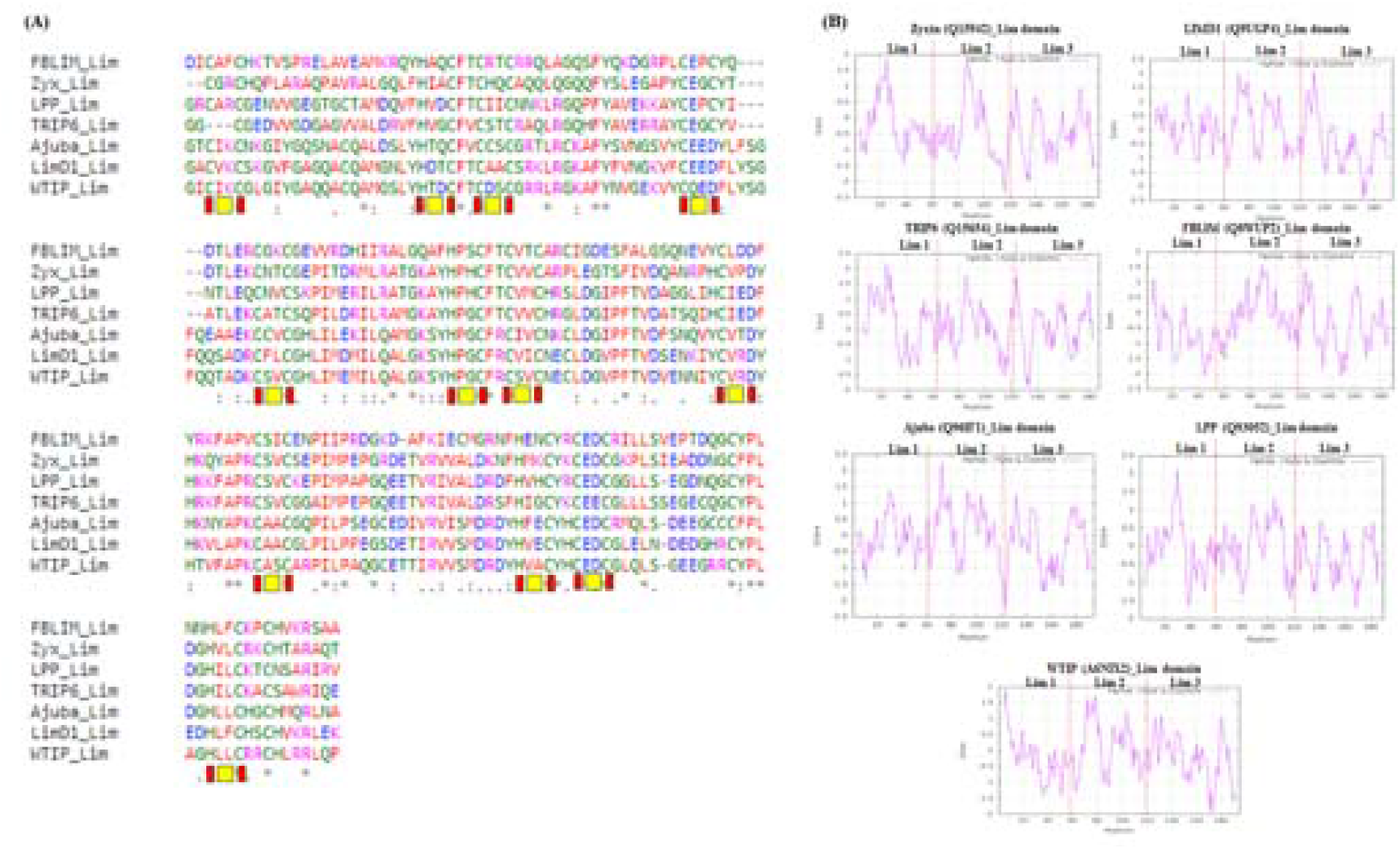
(A) MUSCLE sequence alignment of the Zyxin family Lim domain shows the presence of conserved cysteine/Histidine residues in Lim1/Lim2/Lim3 regions (red boxes are conserved cysteine/histidine/aspartate, Yellow boxes are variable hydrophobic residues) (B) Hydrophobicity plots for Lim domains of Zyxin family proteins. The hydrophobicity analysis was performed using the ProtScale server and plotted against the amino acid position. The scoring from −2.0 to 2.5 on the y-axis reflects hydrophobicity (higher positive (+ve) score depicts more hydrophobicity). This analysis indicates that each Lim domain and Lim1/Lim2/Lim3 region has a distinct hydrophobicity score.

**Supplementary Figure 2:**
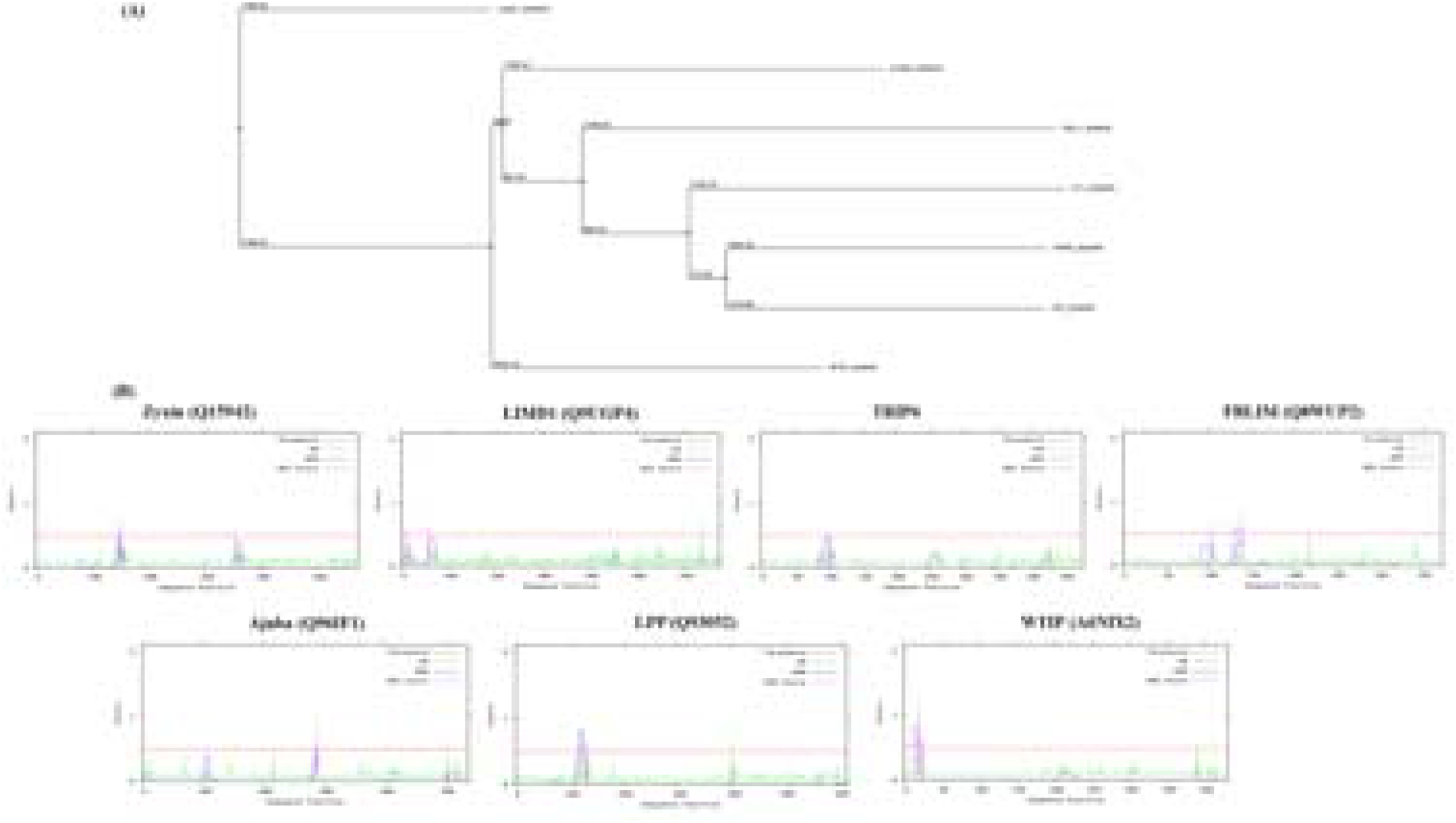
Phylogenetic and NetNES analysis of the Zyxin family proteins. (A) phylogenetic tree created by neighbour joining shows the evolutionary relationship of the zyxin family protein where LIMD1 is most distant member and all other members are more related together (B) NetNES analysis was done to identify the Leucine-rich motif nuclear export sequence (NES) responsible for nuclear transport of protein. The NES was identified through combined analysis implemented by Neural Network (NN), Hidden Markov Model (HMM) and NES scoring algorithm. Only the sequences giving a peak higher than threshold using all three algorithms was identified as NES.

**Supplementary Figure 3:**
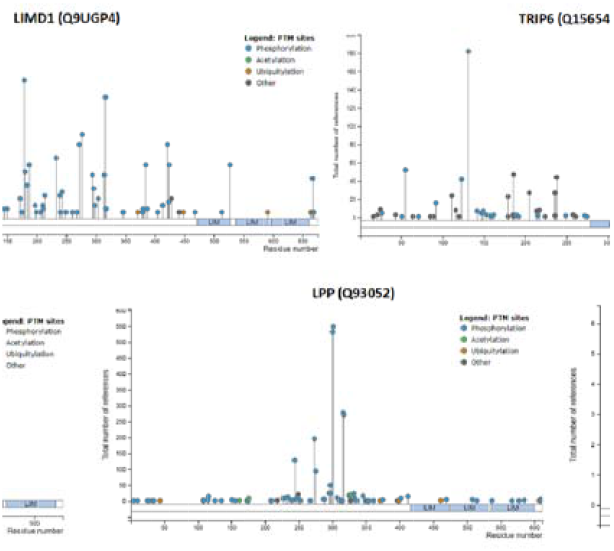
Identification of post-translation modification (PTM) sites of Zyxin family proteins. All proteins exhibited sites for PTM. However, PTMs varied amongst all the member proteins, with the highest PTM observed for LPP, and the lowest for FBLIM1.

**Supplementary Figure 4:**
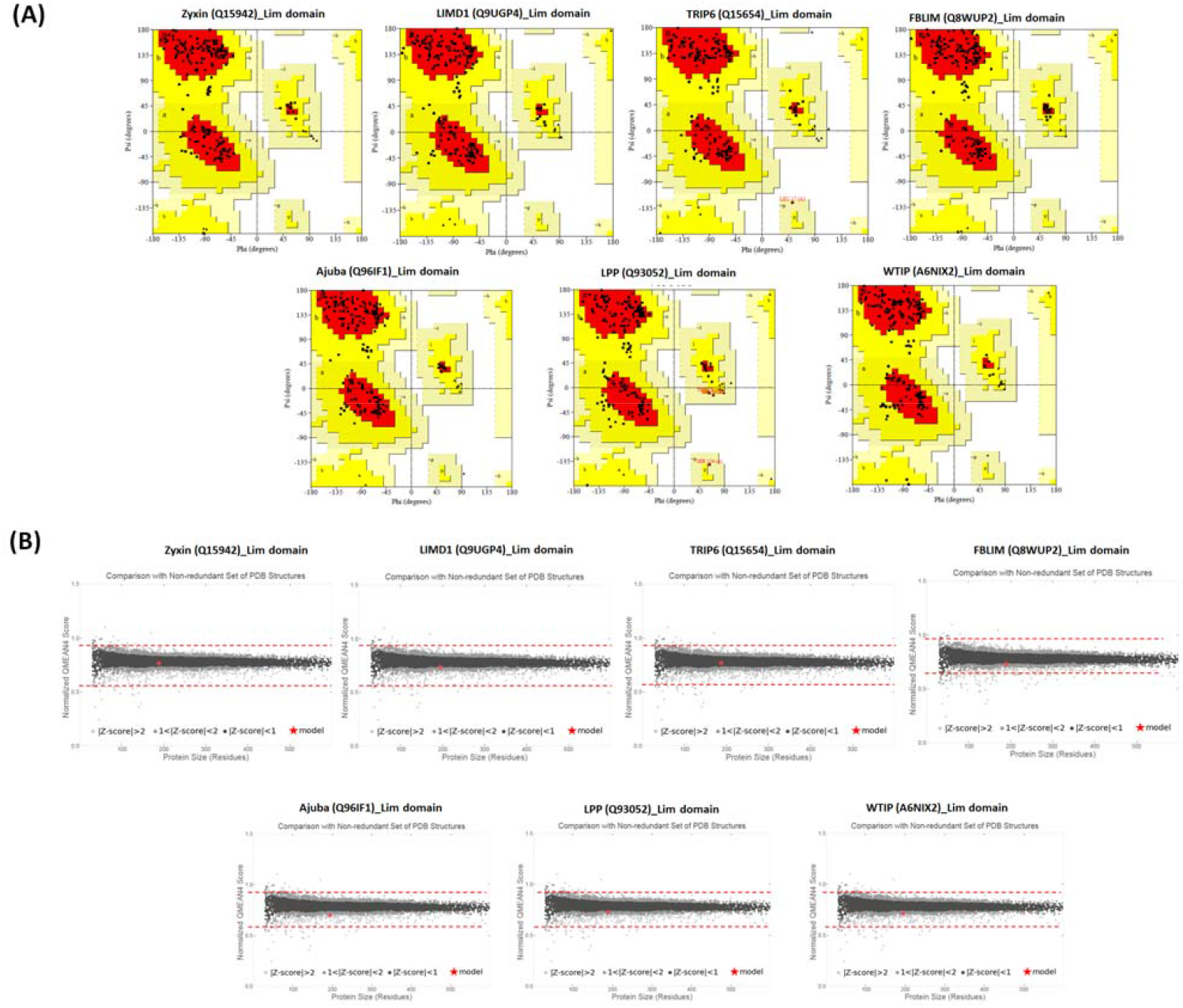
(A) Validation of homology models by Ramachandran plot. Models generated for members of Zyxin family were validated through Ramachandran plots. All models had acceptable Ramachandran values. (B) Absolute quality estimation of all the models by QMEAN, reflecting the native character and quality, red star indicating the query model position while comparing with all available models in PDB.

**Supplementary Figure 5:**
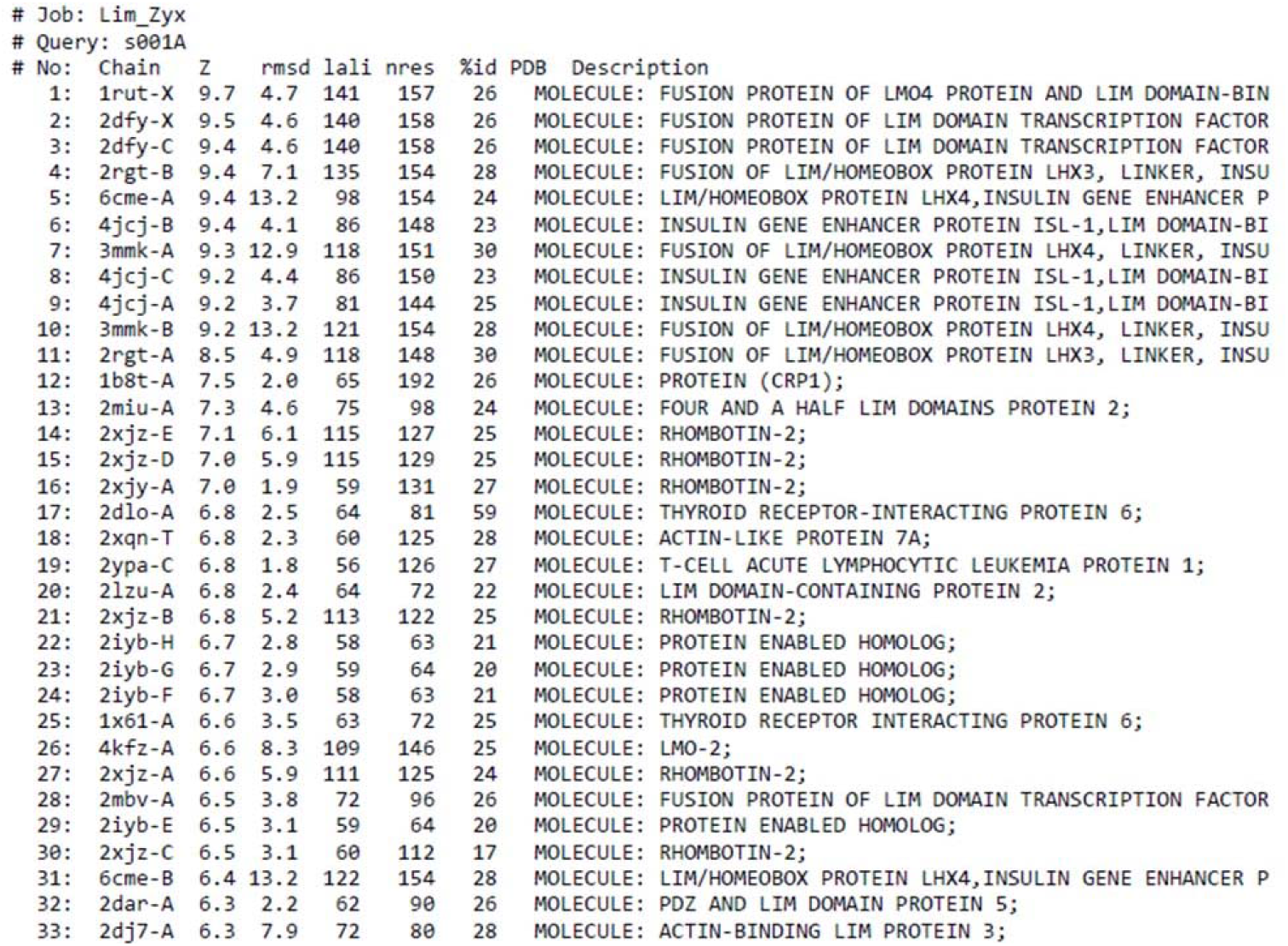
Structural alignment analysis of the Zyxin Lim domain by DALI server, indicates the similarity with transcription regulators.

**Supplementary Figure 6:**
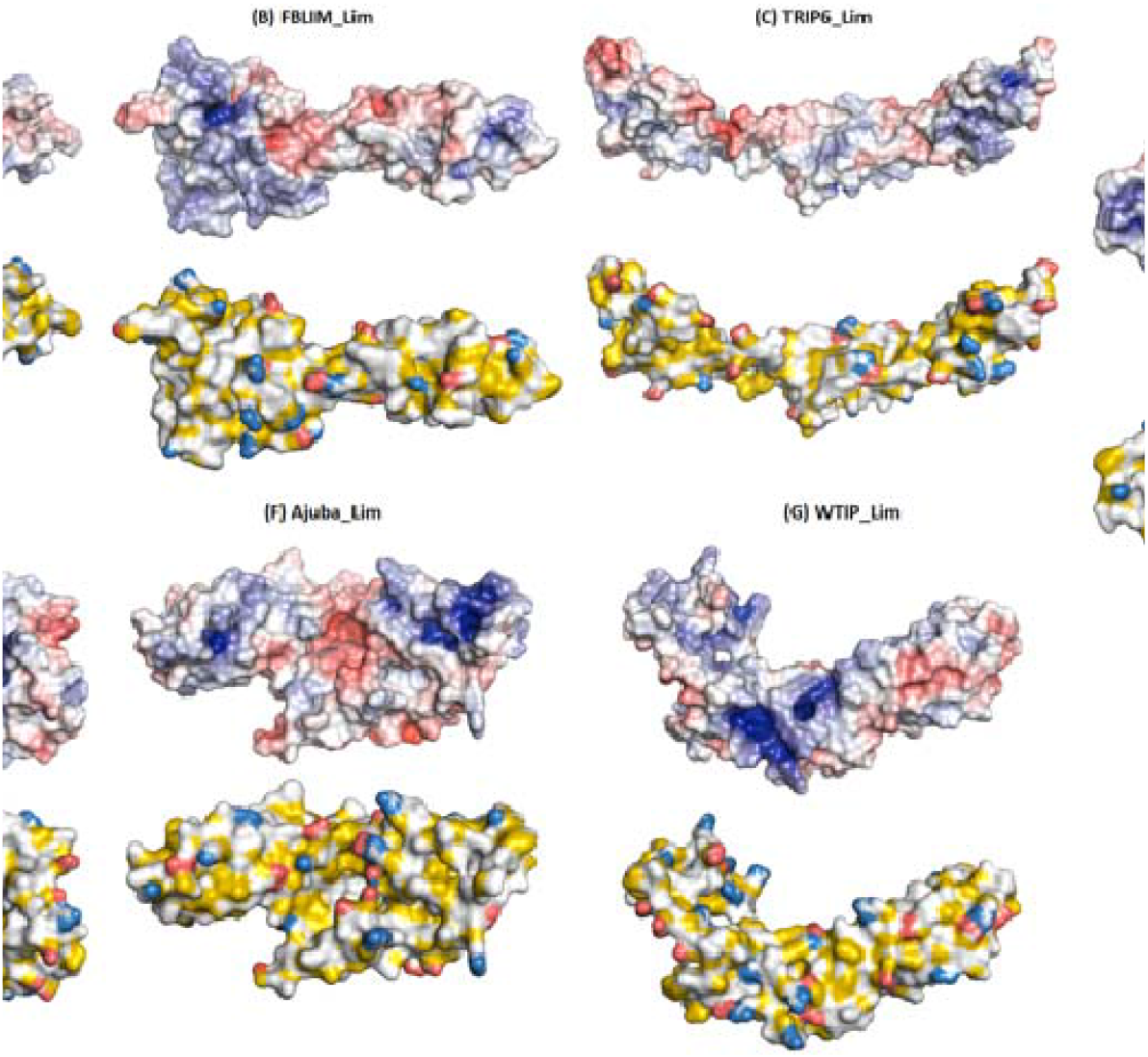
(A) Adaptive Poisson-Boltzmann Solver (APBS) electrostatics and Yellow-Red-Blue (YRB) showing surface potential and hydrophobicity calculations, respectively; of Zyxin family Lim domains.

**Supplementary Figure 7 :**
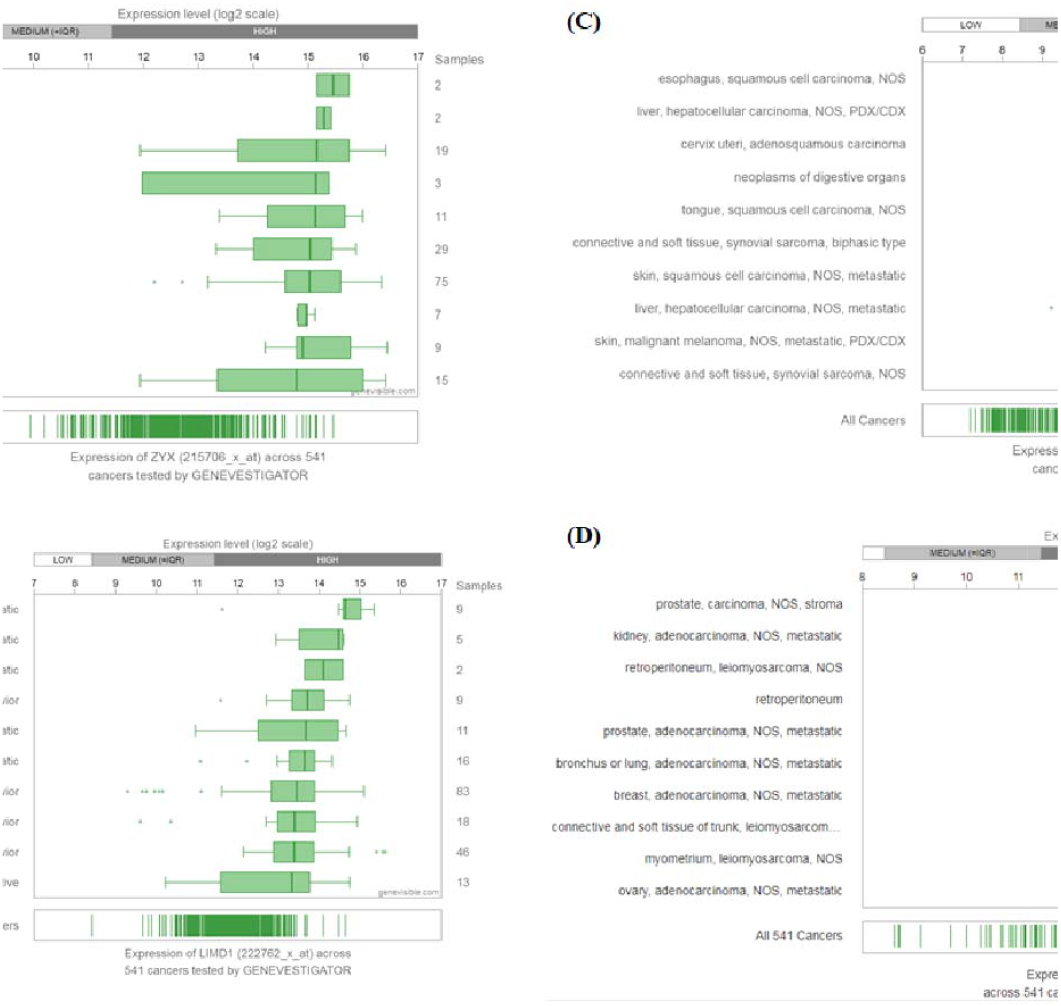
The protein expression profiles of the Zyxin family proteins in different cancers (retrieved from Genevestigator) are presented as bar graphs. (A) Zyxin; (B) LIMD1; (C) Ajuba and (D) LPP. Evidently, the expression of Zyxin family proteins was high in almost all types of cancers.

**Supplementary Figure 8 :**
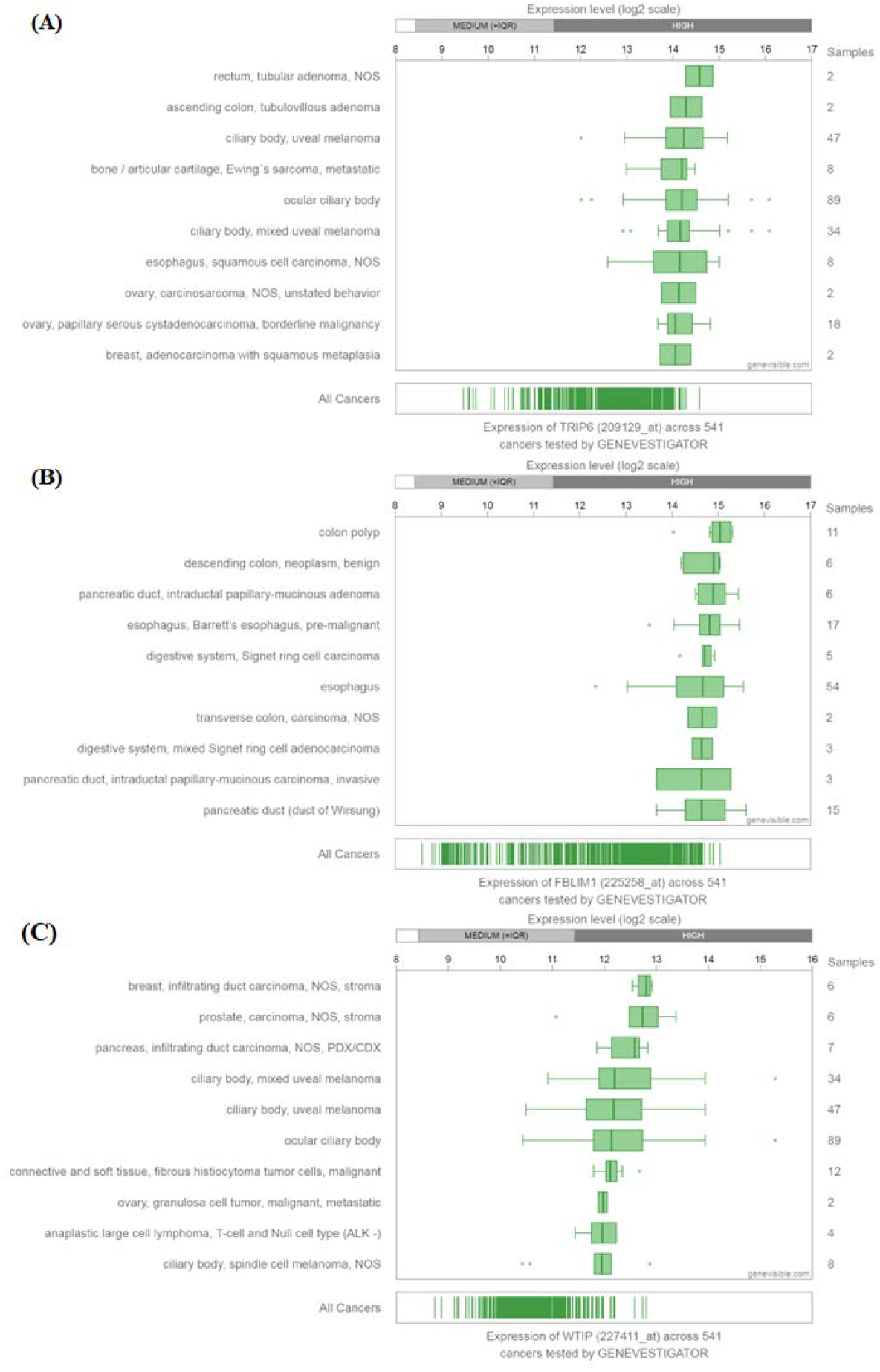
The protein expression profiles of the Zyxin family proteins in different cancers (retrieved from Genevestigator) are presented as bar graphs. (A) TRIP6; (B) FBLIM;; (C) WTIP.. Evidently, the expression of Zyxin family proteins was high in almost all types of cancers.

**Supplementary Figure 9:**
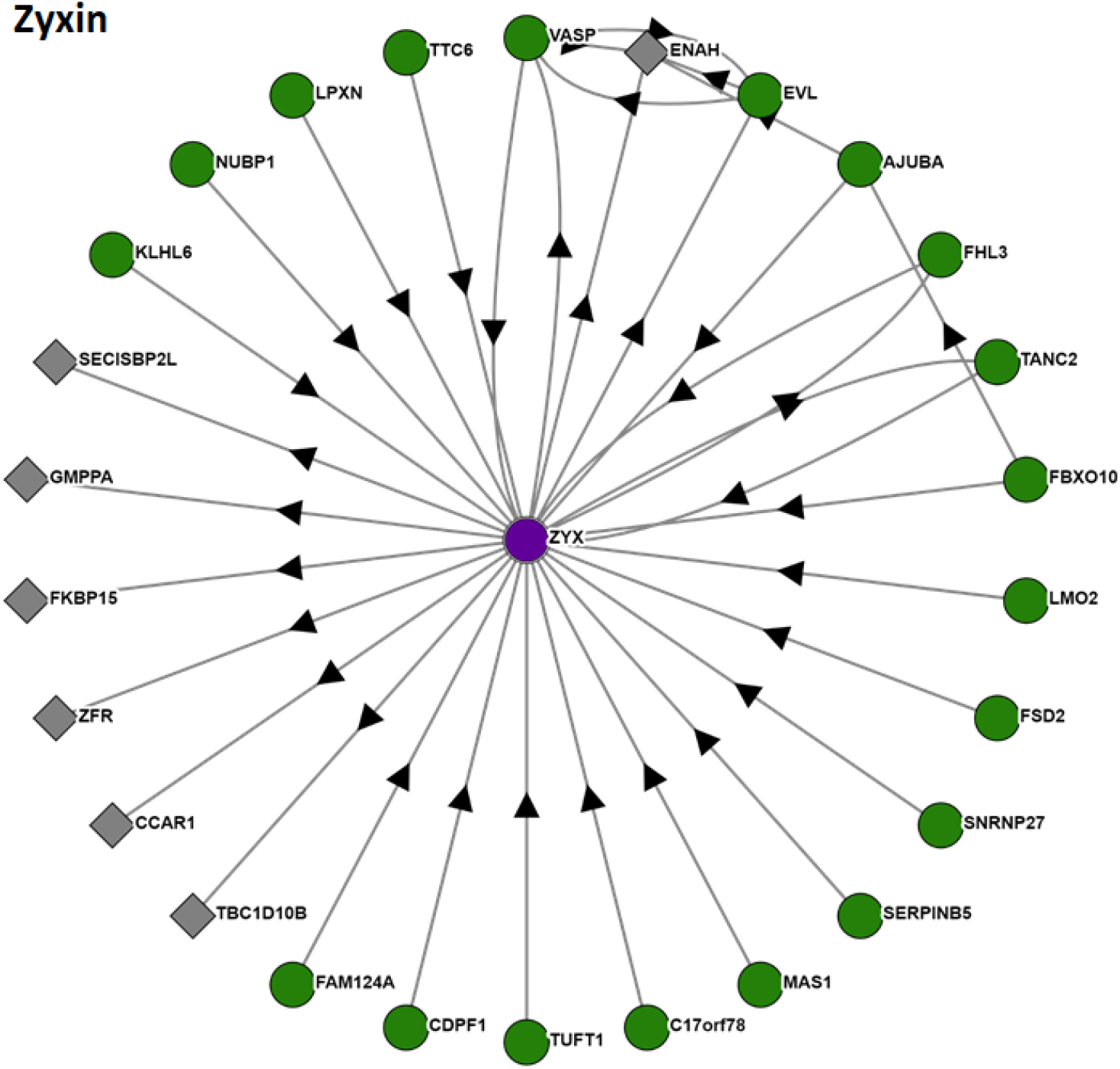
Protein-protein interaction (PPI) analysis of the Zyxin. The arrow indicates the bait and prey relationship of each protein. The bait protein is shown as a green circle whereas the prey protein is shown as grey squares.

**Supplementary Figure 10:**
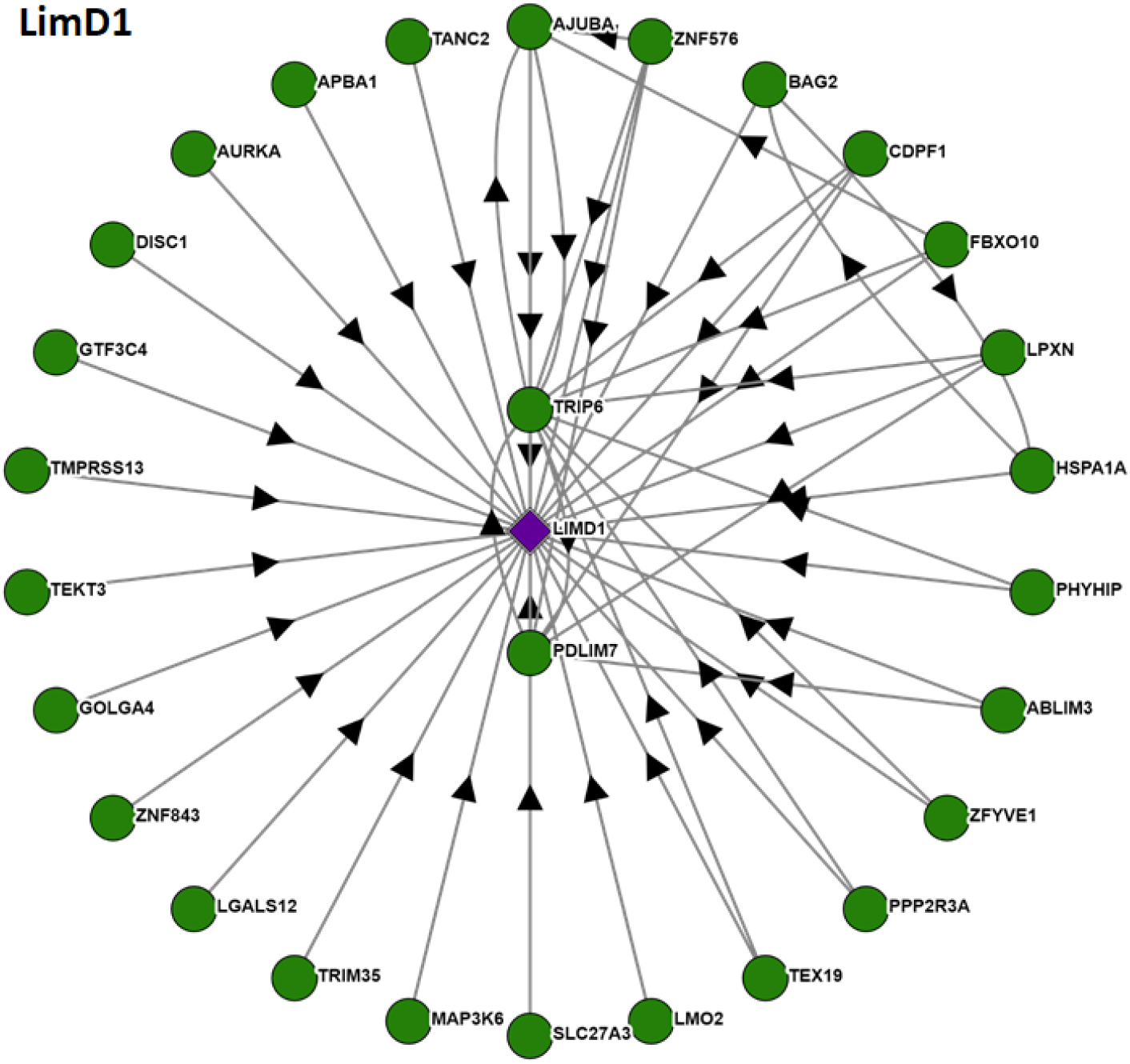
Protein-protein interaction (PPI) analysis of the LimD1. The arrow indicates the bait and prey relationship of each protein. The bait protein is shown as a green circle whereas the prey protein is shown as grey squares.

**Supplementary Figure 11:**
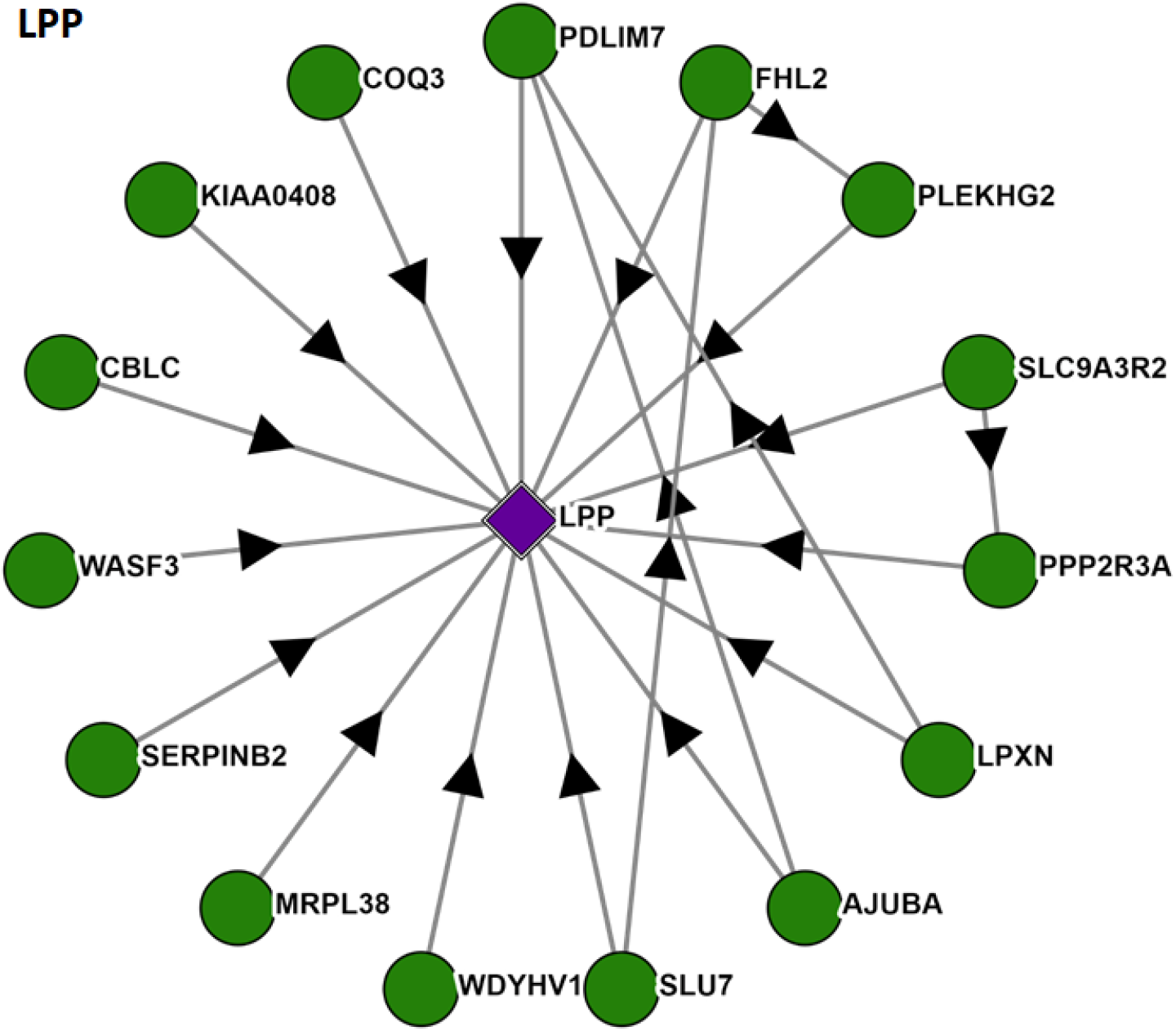
Protein-protein interaction (PPI) analysis of the LPP. The arrow indicates the bait and prey relationship of each protein. The bait protein is shown as a green circle whereas the prey protein is shown as grey squares.

**Supplementary Figure 12:**
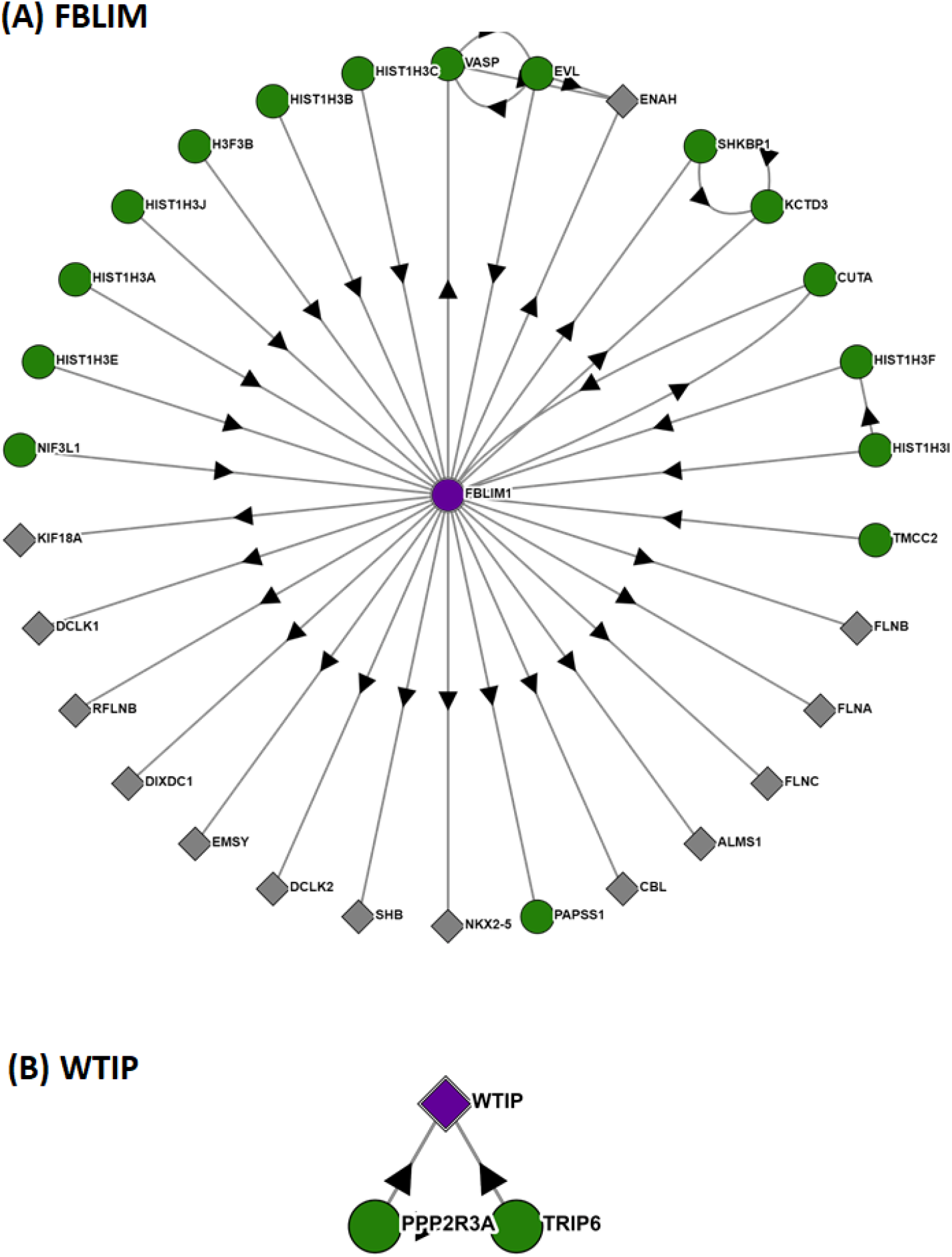
Protein-protein interaction (PPI) analysis of the (A) FBLIM and (B) WTIP. The arrow indicates the bait and prey relationship of each protein. The bait protein is shown as a green circle whereas the prey protein is shown as grey squares. The PPI analysis suggests WTIP has the least interacting partners.

**Supplementary Figure 13:**
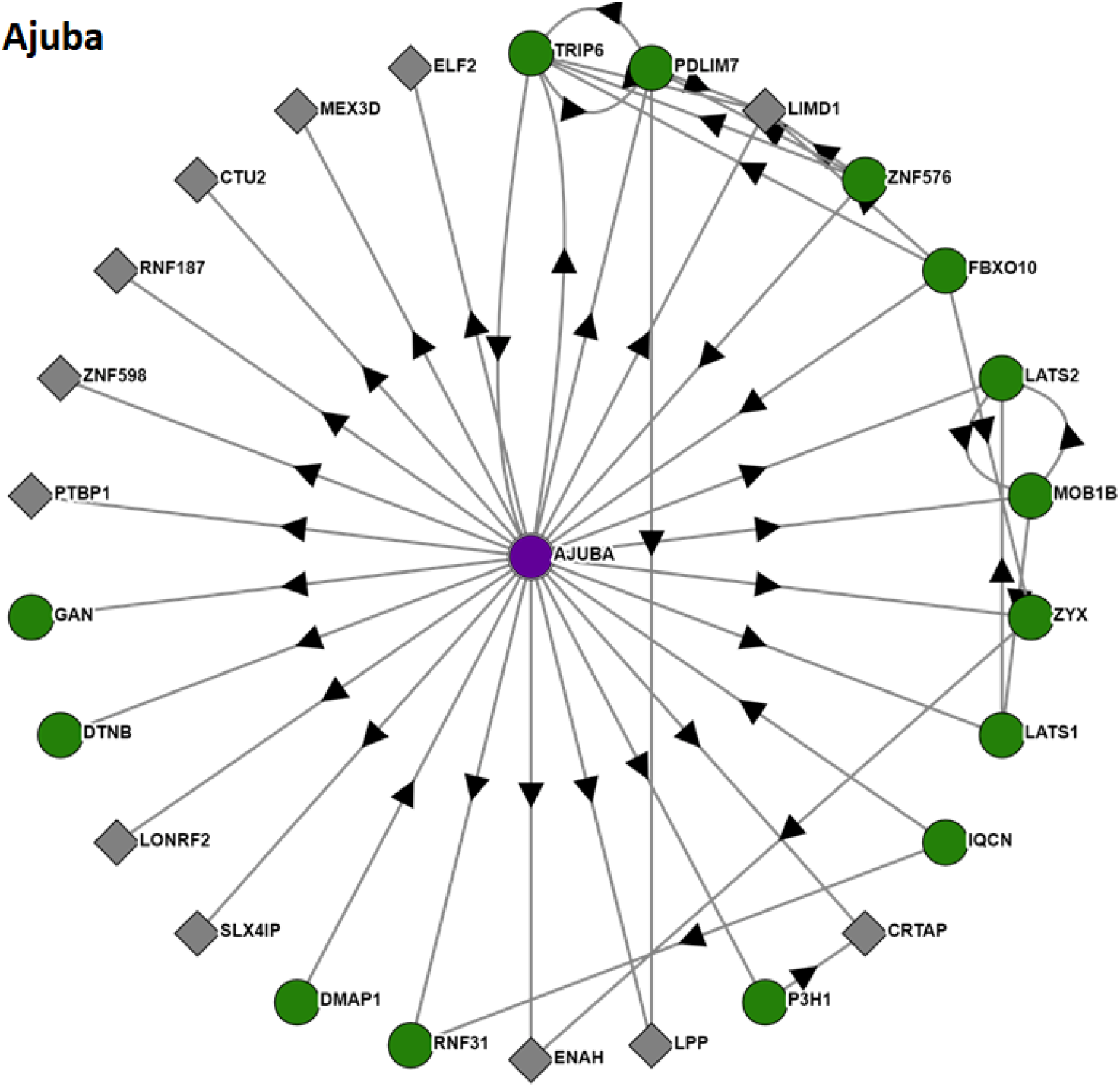
Protein-protein interaction (PPI) analysis of the Ajuba. The arrow indicates the bait and prey relationship of each protein. The bait protein is shown as a green circle whereas the prey protein is shown as grey squares.

**Supplementary Figure 14:**
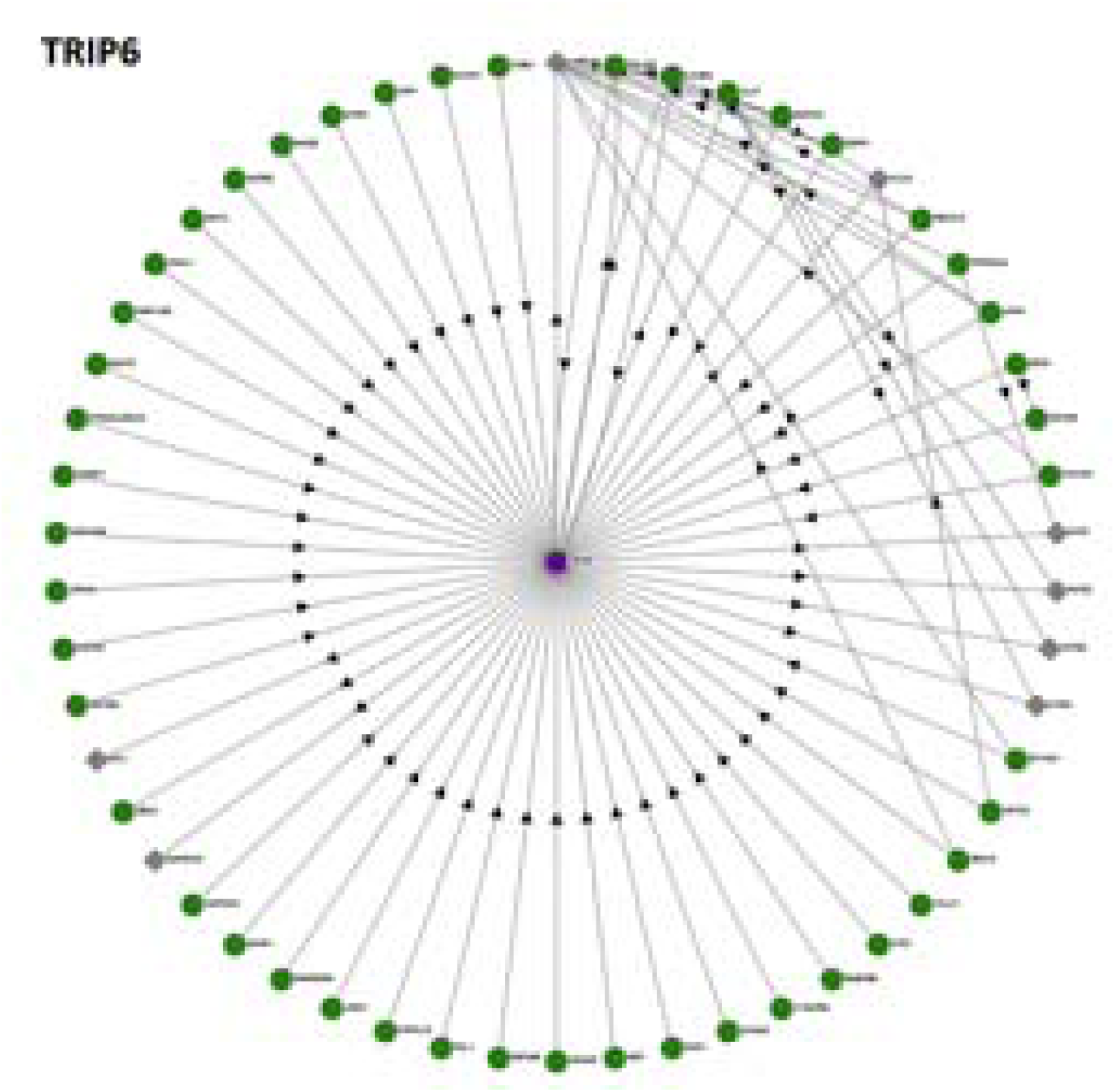
Protein-protein interaction (PPI) analysis of TRIP6. The arrow indicates the bait and prey relationship of each protein. The bait protein is shown as a green circle whereas the prey protein is shown as grey squares. The PPI analysis suggests that TRIP6 has the maximum number of interacting partners.

**Supplementary Figure 15:**
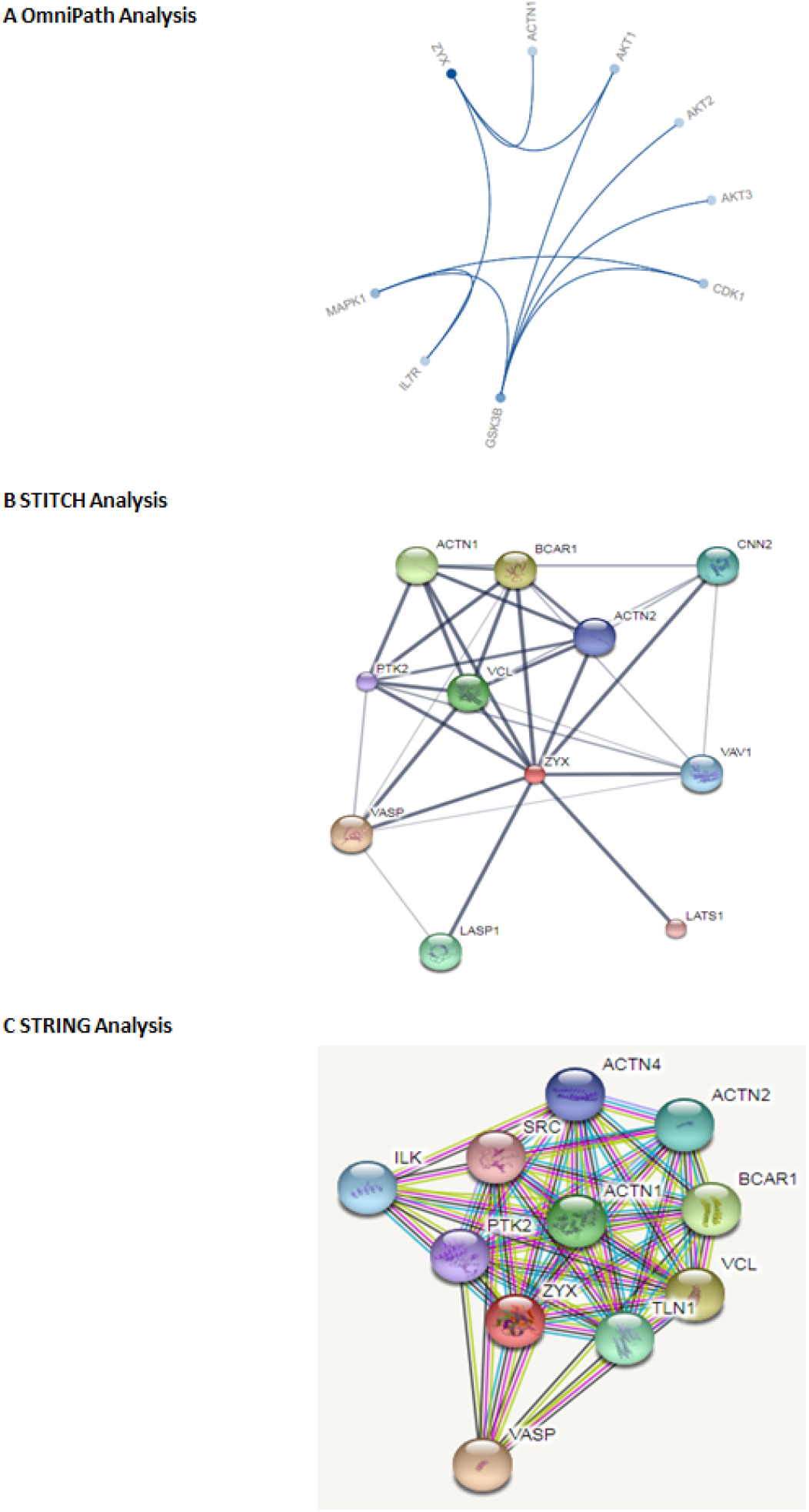
Protein-protein interactions and pathway analysis, (A) Omnipath showing the zyxin involvement in Akt/IL7R pathway; (B) STITCH analysis; (C) STRING analysis.

**Supplementary Table 1:**
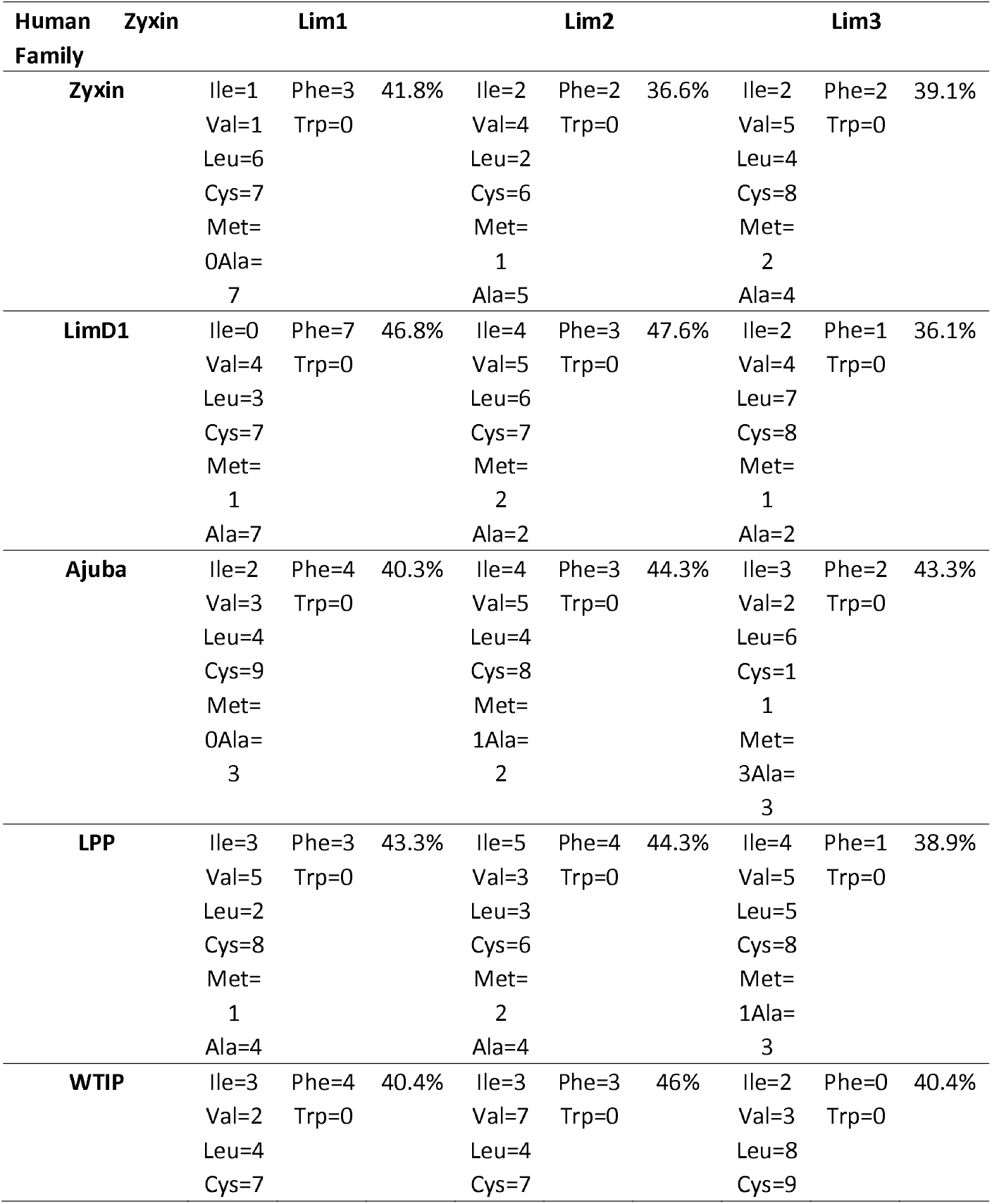
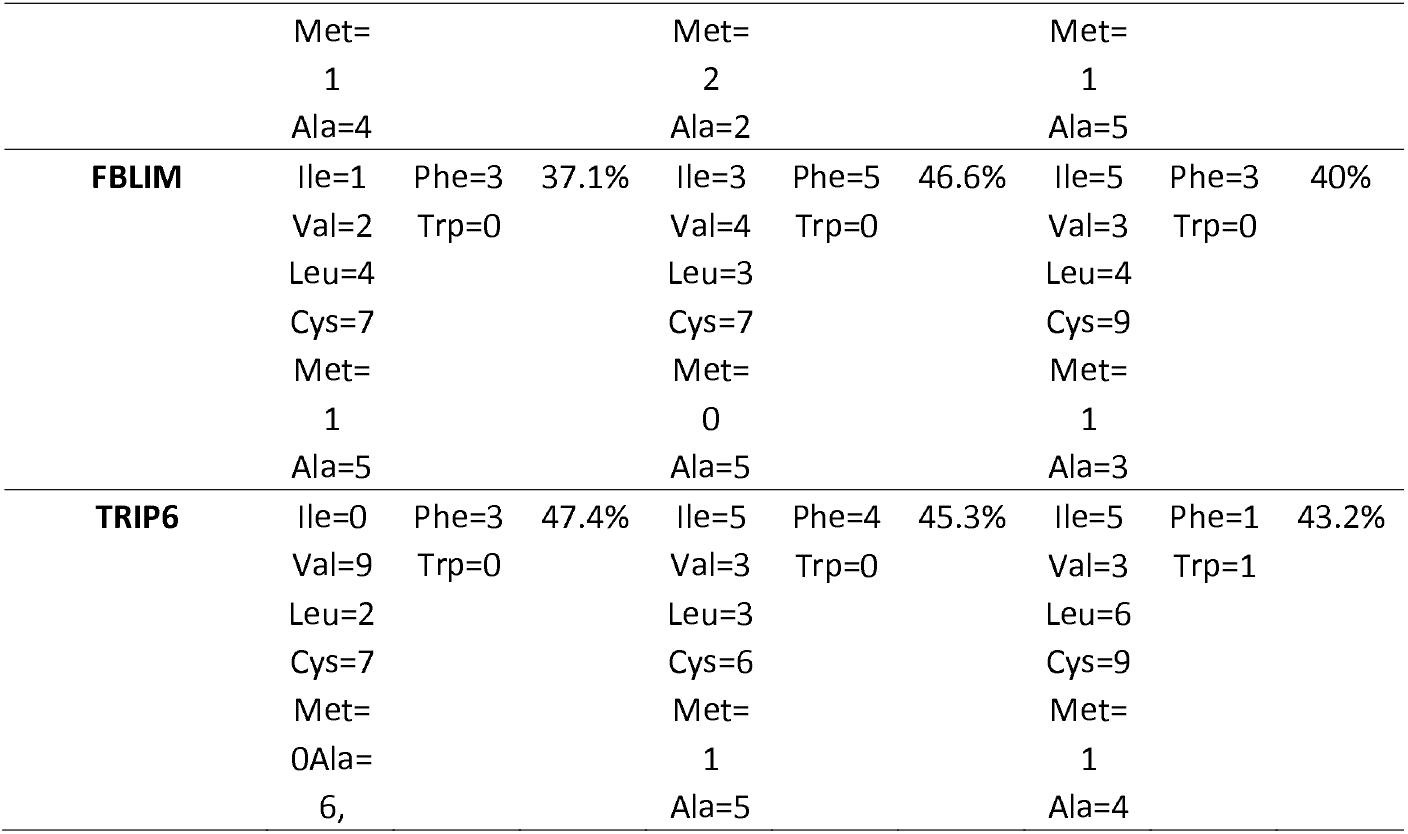
Distribution of hydrophobic amino acids in each LIM domain of human Zyxin family proteins. The column for each LIM domain shows number of hydrophobic amino acids and the total hydrophobicity. These values were determined using Expasy ProtParam, the hydrophobicity of each domain varies from ~36 to 47 %. The number of hydrophobic amino acid varied in each domain but tryptophan was absent amongst all LIMs except LIM3 of TRIP6.

**Supplementary Table 2:** Ohnolog analyis of Zyxin family proteins

| Family Size | Ohnolog Family |
| --- | --- |
| 3 | AJUBA, LIMD1, WTIP |
| 4 | FBLIM1, LPP, TRIP6, ZYX |

**Supplementary Table 3:**
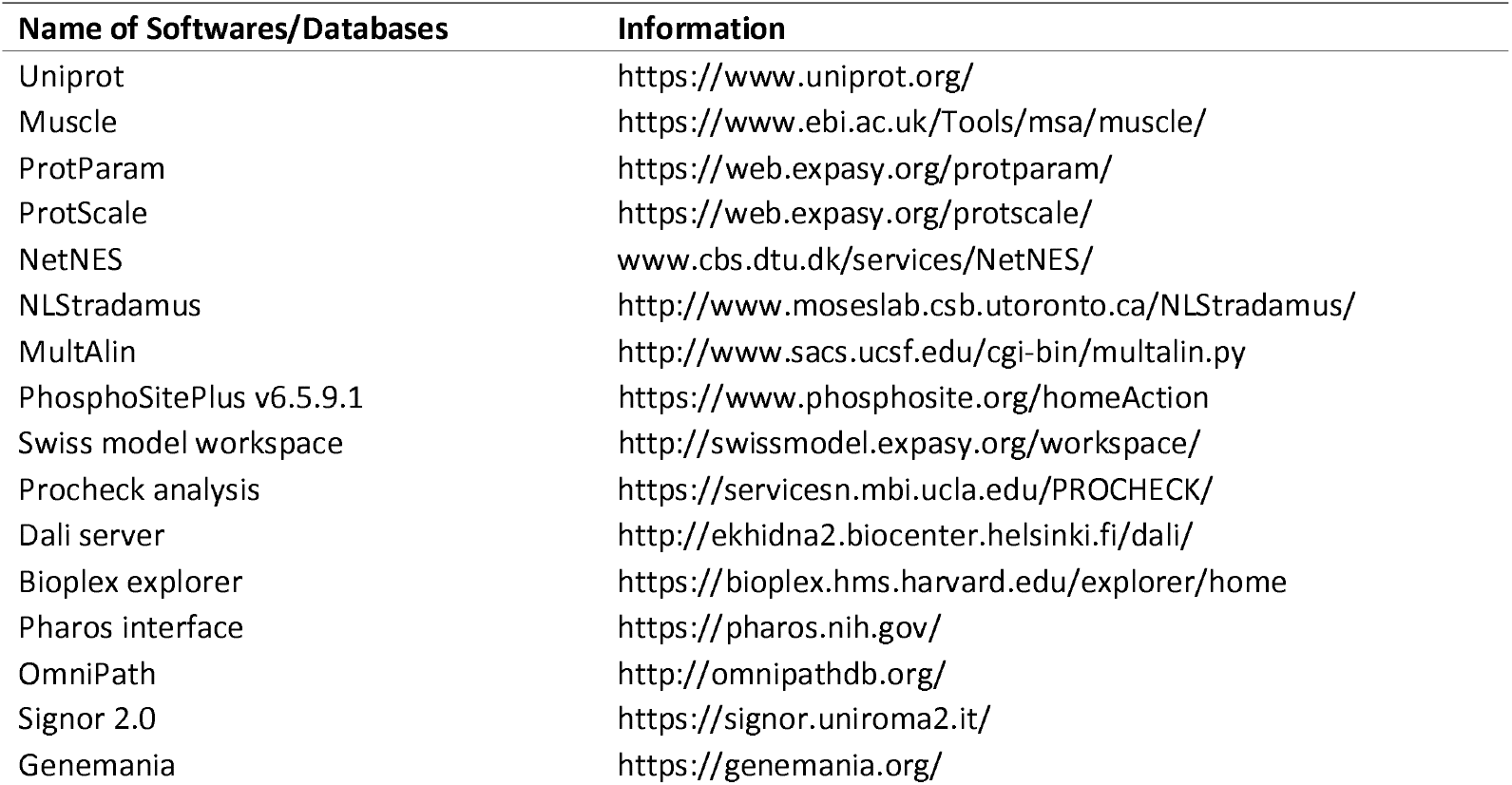
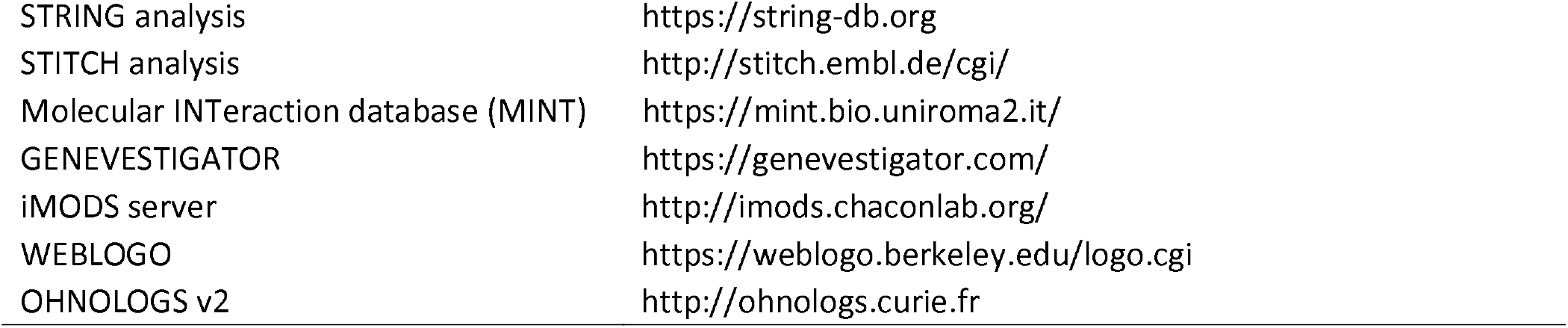
List of all online servers/databases used in the study

| Name of Softwares/Databases | Information |
| --- | --- |
| Uniprot | https://www.uniprot.org/ |
| Muscle | https://www.ebi.ac.uk/Tools/msa/muscle/ |
| ProtParam | https://web.expasy.org/protparam/ |
| ProtScale | https://web.expasy.org/protscale/ |
| NetNES | www.cbs.dtu.dk/services/NetNES/ |
| NLStradamus | http://www.moseslab.csb.utoronto.ca/NLStradamus/ |
| MultAlin | http://www.sacs.ucsf.edu/cgi-bin/multalin.py |
| PhosphoSitePlus v6.5.9.1 | https://www.phosphosite.org/homeAction |
| Swiss model workspace | http://swissmodel.expasy.org/workspace/ |
| Procheck analysis | https://servicesn.mbi.ucla.edu/PROCHECK/ |
| Dali server | http://ekhidna2.biocenter.helsinki.fi/dali/ |
| Bioplex explorer | https://bioplex.hms.harvard.edu/explorer/home |
| Pharos interface | https://pharos.nih.gov/ |
| OmniPath | http://omnipathdb.org/ |
| Signor 2.0 | https://signor.uniroma2.it/ |
| Genemania | https://genemania.org/ |
| STRING analysis | https://string-db.org |
| STITCH analysis | http://stitch.embl.de/cgi/ |
| Molecular INTERaction database (MINT) | https://mint.bio.uniroma2.it/ |
| GENEVESTIGATOR | https://genevestigator.com/ |
| iMODS server | http://imods.chaconlab.org/ |
| WEBLOGO | https://weblogo.berkeley.edu/logo.cgi |
| OHNOLOGS v2 | http://ohnologs.curie.fr |

## Author Contributions

Conceptualization: M.Q.S. and T.R.P.; Methodology: M.Q.S.; Validation, M.Q.S., M.D.B.,T.R.P.; Formal Analysis, M.Q.S. and M.D.B.; Investigation, M.Q.S. and M.D.B.; Writing and Original Draft Preparation, M.Q.S. and M.D.B.; Writing – Review & Editing, M.Q.S., M.D.B. and T.R.P.; Supervision, T.R.P; Project Administration, T.R.P.; Funding Acquisition, T.R.P. All authors have read and agreed to the published version of the manuscript.

## References

1. Kotb, A., M.E. Hyndman, and T.R. Patel, The role of zyxin in regulation of malignancies. Heliyon, 2018. 4(7): p. e00695.

2. van der Gaag, E.J., et al., Role of zyxin in differential cell spreading and proliferation of melanoma cells and melanocytes. J Invest Dermatol, 2002. 118(2): p. 246–54.

3. Zhou, J., et al., Zyxin promotes colon cancer tumorigenesis in a mitotic phosphorylation-dependent manner and through CDK8-mediated YAP activation. Proc Natl Acad Sci U S A, 2018. 115(29): p. E6760–E6769.

4. Gaspar, P., et al., Zyxin antagonizes the FERM protein expanded to couple F-actin and Yorkie-dependent organ growth. Curr Biol, 2015. 25(6): p. 679–689.

5. Rauskolb, C., et al., Zyxin links fat signaling to the hippo pathway. PLoS Biol, 2011. 9(6): p. e1000624.

6. Sy, S.M., et al., Novel identification of zyxin upregulations in the motile phenotype of hepatocellular carcinoma. Mod Pathol, 2006. 19(8): p. 1108–16.

7. Mise, N., et al., Zyxin is a transforming growth factor-beta (TGF-beta)/Smad3 target gene that regulates lung cancer cell motility via integrin alpha5beta1. J Biol Chem, 2012. 287(37): p. 31393–405.

8. Zhong, C., et al., Zyxin as a potential cancer prognostic marker promotes the proliferation and metastasis of colorectal cancer cells. J Cell Physiol, 2019.

9. Neuwirt, H., et al., Cancer-associated fibroblasts promote prostate tumor growth and progression through upregulation of cholesterol and steroid biosynthesis. Cell Commun Signal, 2020. 18(1): p. 11.

10. Wen, X.M., et al., Zyxin (ZYX) promotes invasion and acts as a biomarker for aggressive phenotypes of human glioblastoma multiforme. Lab Invest, 2020.

11. Wu, Z., et al., WT1-interacting protein inhibits cell proliferation and tumorigenicity in non-small-cell lung cancer via the AKT/FOXO1 axis. Mol Oncol, 2019. 13(5): p. 1059–1074.

12. James, V., et al., LIM-domain proteins, LIMD1, Ajuba, and WTIP are required for microRNA-mediated gene silencing. Proc Natl Acad Sci U S A, 2010. 107(28): p. 12499–504.

13. Ou, Y., et al., Migfilin protein promotes migration and invasion in human glioma through epidermal growth factor receptor-mediated phospholipase C-gamma and STAT3 protein signaling pathways. J Biol Chem, 2012. 287(39): p. 32394–405.

14. Kiepas, A., et al., The SHCA adapter protein cooperates with lipoma-preferred partner in the regulation of adhesion dynamics and invadopodia formation. J Biol Chem, 2020.

15. Kuriyama, S., et al., LPP inhibits collective cell migration during lung cancer dissemination. Oncogene, 2016. 35(8): p. 952–64.

16. Feuerstein, R., et al., The LIM/double zinc-finger motif functions as a protein dimerization domain. Proc Natl Acad Sci U S A, 1994. 91(22): p. 10655–9.

17. Schmeichel, K.L. and M.C. Beckerle, The LIM domain is a modular protein-binding interface. Cell, 1994. 79(2): p. 211–9.

18. Kadrmas, J.L. and M.C. Beckerle, The LIM domain: from the cytoskeleton to the nucleus. Nat Rev Mol Cell Biol, 2004. 5(11): p. 920–31.

19. Michelsen, J.W., et al., The LIM motif defines a specific zinc-binding protein domain. Proc Natl Acad Sci U S A, 1993. 90(10): p. 4404–8.

20. Minor, D.L., Jr. and P.S. Kim, Measurement of the beta-sheet-forming propensities of amino acids. Nature, 1994. 367(6464): p. 660–3.

21. Singh, P.P. and H. Isambert, OHNOLOGS v2: a comprehensive resource for the genes retained from whole genome duplication in vertebrates. Nucleic Acids Res, 2020. 48(D1): p. D724–D730.

22. la Cour, T., et al., Analysis and prediction of leucine-rich nuclear export signals. Protein Eng Des Sel, 2004. 17(6): p. 527–36.

23. Kiraga, J., et al., The relationships between the isoelectric point and: length of proteins, taxonomy and ecology of organisms. BMC Genomics, 2007. 8: p. 163.

24. Schwartz, R., C.S. Ting, and J. King, Whole proteome pI values correlate with subcellular localizations of proteins for organisms within the three domains of life. Genome Res, 2001. 11(5): p. 703–9.

25. Zhu, K., et al., Protein pI shifts due to posttranslational modifications in the separation and characterization of proteins. Anal Chem, 2005. 77(9): p. 2745–55.

26. Hornbeck, P.V., et al., 15 years of PhosphoSitePlus(R): integrating post-translationally modified sites, disease variants and isoforms. Nucleic Acids Res, 2019. 47(D1): p. D433–D441.

27. Petit, M.M., et al., LPP, an actin cytoskeleton protein related to zyxin, harbors a nuclear export signal and transcriptional activation capacity. Mol Biol Cell, 2000. 11(1): p. 117–29.

28. Laskowski, R.A., et al., PROCHECK: a program to check the stereochemical quality of protein structures. Journal of applied crystallography, 1993. 26(2): p. 283–291.

29. Benkert, P., M. Biasini, and T. Schwede, Toward the estimation of the absolute quality of individual protein structure models. Bioinformatics, 2011. 27(3): p. 343–50.

30. Castello, A., et al., Insights into RNA biology from an atlas of mammalian mRNA-binding proteins. Cell, 2012. 149(6): p. 1393–406.

31. Kumar, M., M.M. Gromiha, and G.P. Raghava, Identification of DNA-binding proteins using support vector machines and evolutionary profiles. BMC Bioinformatics, 2007. 8: p. 463.

32. Holm, L. and P. Rosenstrom, Dali server: conservation mapping in 3D. Nucleic Acids Res, 2010. 38(Web Server issue): p. W545–9.

33. Sam, V., et al., ROC and confusion analysis of structure comparison methods identify the main causes of divergence from manual protein classification. BMC Bioinformatics, 2006. 7: p. 206.

34. Deane, J.E., et al., Tandem LIM domains provide synergistic binding in the LMO4:Ldb1 complex. EMBO J, 2004. 23(18): p. 3589–98.

35. Jeffries, C.M., et al., Stabilization of a binary protein complex by intein-mediated cyclization. Protein Sci, 2006. 15(11): p. 2612–8.

36. Bhati, M., et al., Implementing the LIM code: the structural basis for cell type-specific assembly of LIM-homeodomain complexes. EMBO J, 2008. 27(14): p. 2018–29.

37. Stokes, P.H., et al., Mutation in a flexible linker modulates binding affinity for modular complexes. Proteins, 2019. 87(5): p. 425–429.

38. Gadd, M.S., et al., A structural basis for the regulation of the LIM-homeodomain protein islet 1 (Isl1) by intra-and intermolecular interactions. J Biol Chem, 2013. 288(30):p. 21924–35.

39. Racevskis, J., et al., Molecular cloning of LMO41, a new human LIM domain gene. Biochim Biophys Acta, 1999. 1445(1):p. 148–53.

40. Vu, D., et al., Transcription regulator LMO4 interferes with neuritogenesis in human SH-SY5Y neuroblastoma cells. Brain Res Mol Brain Res, 2003. 115(2):p. 93–103.

41. Sum, E.Y., et al., The LIM domain protein LMO4 interacts with the cofactor CtIP and the tumor suppressor BRCA1 and inhibits BRCA1 activity. J Biol Chem, 2002. 277(10):p. 7849–56.

42. Lopez-Blanco, J.R., et al., iMODS: internal coordinates normal mode analysis server. Nucleic Acids Res, 2014. 42(Web Server issue): p. W271–6.

43. Amadei, A., A.B. Linssen, and H.J. Berendsen, Essential dynamics of proteins. Proteins, 1993. 17(4):p. 412–25.

44. Schweppe, D.K., et al., BioPlex Display: An Interactive Suite for Large-Scale AP-MS Protein-Protein Interaction Data. J Proteome Res, 2018. 17(1): p. 722–726.

45. Warde-Farley, D., et al., The GeneMANIA prediction server: biological network integration for gene prioritization and predicting gene function. Nucleic Acids Res, 2010. 38(Web Server issue): p. W214–20.

46. Licata, L., et al., MINT, the molecular interaction database: 2012 update. Nucleic Acids Res, 2012. 40(Database issue): p. D857–61.

47. Szklarczyk, D., et al., STITCH 5: augmenting protein-chemical interaction networks with tissue and affinity data. Nucleic Acids Res, 2016. 44(D1): p. D380–4.

48. Licata, L., et al., SIGNOR 2.0, the SIGnaling Network Open Resource 2.0: 2019 update. Nucleic Acids Res, 2020. 48(D1): p. D504–D510.

49. Szklarczyk, D., et al., STRING v11: protein-protein association networks with increased coverage, supporting functional discovery in genome-wide experimental datasets. Nucleic Acids Res, 2019. 47(D1): p. D607–D613.

50. Bailey, M.H., et al., Comprehensive Characterization of Cancer Driver Genes and Mutations. Cell, 2018. 173(2):p. 371–385 e18.

51. Chen, H.K., et al., Human Nopp140, which interacts with RNA polymerase I: implications for rRNA gene transcription and nucleolar structural organization. Mol Cell Biol, 1999. 19(12): p. 8536–46.

52. Chan, C.B., et al., Akt phosphorylation of zyxin mediates its interaction with acinus-S and prevents acinus-triggered chromatin condensation. Cell Death Differ, 2007. 14(9): p. 1688–99.

53. Abassi, Y.A., et al., p130Cas Couples the tyrosine kinase Bmx/Etk with regulation of the actin cytoskeleton and cell migration. J Biol Chem, 2003. 278(37): p. 35636–43.

54. Sakakibara, A., et al., Novel function of Chat in controlling cell adhesion via Cas-Crk-C3G-pathway-mediated Rap1 activation. J Cell Sci, 2002. 115(Pt 24): p. 4915–24.

55. de Jesus, A.J. and T.W. Allen, The role of tryptophan side chains in membrane protein anchoring and hydrophobic mismatch. Biochim Biophys Acta, 2013. 1828(2): p. 864–76.

56. Samanta, U. and P. Chakrabarti, Assessing the role of tryptophan residues in the binding site. Protein Eng, 2001. 14(1): p. 7–15.

57. Makino, T. and A. McLysaght, Ohnologs in the human genome are dosage balanced and frequently associated with disease. Proc Natl Acad Sci U S A, 2010. 107(20): p. 9270–4.

58. Koch, B.J., J.F. Ryan, and A.D. Baxevanis, The diversification of the LIM superclass at the base of the metazoa increased subcellular complexity and promoted multicellular specialization. PLoS One, 2012. 7(3): p. e33261.

59. Ayyanathan, K., et al., The Ajuba LIM domain protein is a corepressor for SNAG domain mediated repression and participates in nucleocytoplasmic Shuttling. Cancer Res, 2007. 67(19): p. 9097–106.

60. Kim, S.H., et al., The LIM-only transcription factor LMO2 determines tumorigenic and angiogenic traits in glioma stem cells. Cell Death Differ, 2015. 22(9):p. 1517–25.

61. Marie, H., et al., The LIM protein Ajuba is recruited to cadherin-dependent cell junctions through an association with alpha-catenin. J Biol Chem, 2003. 278(2):p. 1220–8.

62. Sanchez-Garcia, I. and T.H. Rabbitts, The LIM domain: a new structural motif found in zinc-finger-like proteins. Trends Genet, 1994. 10(9): p. 315–20.

63. Wu, R., et al., Specificity of LIM domain interactions with receptor tyrosine kinases. J Biol Chem, 1996. 271(27): p. 15934–41.

64. Kamberaj, H. and A. van der Vaart, Extracting the causality of correlated motions from molecular dynamics simulations. Biophys J, 2009. 97(6): p. 1747–55.

65. Goodey, N.M. and S.J. Benkovic, Allosteric regulation and catalysis emerge via a common route. Nat Chem Biol, 2008. 4(8): p. 474–82.

66. Hammes, G.G., Multiple conformational changes in enzyme catalysis. Biochemistry, 2002. 41(26): p. 8221–8.

67. Jarymowycz, V.A. and M.J. Stone, Fast time scale dynamics of protein backbones: NMR relaxation methods, applications, and functional consequences. Chem Rev, 2006. 106(5):p. 1624–71.

68. Magrane, M. and C. UniProt, UniProt Knowledgebase: a hub of integrated protein data. Database (Oxford), 2011. 2011: p. bar009.

69. Edgar, R.C., MUSCLE: a multiple sequence alignment method with reduced time and space complexity. BMC Bioinformatics, 2004. 5: p. 113.

70. Thompson, J.D., T.J. Gibson, and D.G. Higgins, Multiple sequence alignment using ClustalW and ClustalX. Curr Protoc Bioinformatics, 2002. Chapter 2:p. Unit 2 3.

71. Waterhouse, A.M., et al., Jalview Version 2--a multiple sequence alignment editor and analysis workbench. Bioinformatics, 2009. 25(9): p. 1189–91.

72. Eddy, S.R., Where did the BLOSUM62 alignment score matrix come from? Nat Biotechnol, 2004. 22(8):p. 1035–6.

73. Edgar, R.C., MUSCLE: multiple sequence alignment with high accuracy and high throughput. Nucleic Acids Res, 2004. 32(5):p. 1792–7.

74. Crooks, G.E., et al., WebLogo: a sequence logo generator. Genome Res, 2004. 14(6): p. 1188–90.

75. Nguyen Ba, A.N., et al., NLStradamus: a simple Hidden Markov Model for nuclear localization signal prediction. BMC Bioinformatics, 2009. 10:p. 202.

76. Corpet, F., Multiple sequence alignment with hierarchical clustering. Nucleic Acids Res, 1988. 16(22): p. 10881–90.

77. Artimo, P., et al., ExPASy: SIB bioinformatics resource portal. Nucleic Acids Res, 2012. 40(Web Server issue): p. W597–603.

78. Kim, D.E., D. Chivian, and D. Baker, Protein structure prediction and analysis using the Robetta server. Nucleic acids research, 2004. 32(suppl_2): p. W526–W531.

79. Kim, D.E., D. Chivian, and D. Baker, Protein structure prediction and analysis using the Robetta server. Nucleic Acids Res, 2004. 32(Web Server issue): p. W526–31.

80. Arnold, K., et al., The SWISS-MODEL workspace: a web-based environment for protein structure homology modelling. Bioinformatics, 2006. 22(2):p. 195–201.

81. Bordoli, L. and T. Schwede, Automated protein structure modeling with SWISS-MODEL Workspace and the Protein Model Portal. Methods Mol Biol, 2012. 857:p. 107–36.

82. Holm, L. and L.M. Laakso, Dali server update. Nucleic Acids Res, 2016. 44(W1): p. W351–5.

83. Baker, N.A., et al., Electrostatics of nanosystems: application to microtubules and the ribosome. Proc Natl Acad Sci U S A, 2001. 98(18): p. 10037–41.

84. Lopez-Blanco, J.R., et al., Exploring large macromolecular functional motions on clusters of multicore processors. Journal of Computational Physics, 2013. 246:p. 275–288.

85. Nguyen, D.T., et al., Pharos: Collating protein information to shed light on the druggable genome. Nucleic Acids Res, 2017. 45(D1): p. D995–D1002.

86. Turei, D., T. Korcsmaros, and J. Saez-Rodriguez, OmniPath: guidelines and gateway for literature-curated signaling pathway resources. Nat Methods, 2016. 13(12): p. 966–967.

87. Hruz, T., et al., Genevestigator v3: a reference expression database for the meta-analysis of transcriptomes. Adv Bioinformatics, 2008. 2008: p. 420747.

